# The macroecology and evolution of avian competence for *Borrelia burgdorferi*

**DOI:** 10.1101/2020.04.15.040352

**Authors:** Daniel J. Becker, Barbara A. Han

## Abstract

**Aim:** Predicting novel reservoirs of zoonotic pathogens would be improved by identifying inter-specific drivers of host competence, the ability to transmit pathogens to new hosts or vectors. Tick-borne pathogens can provide a useful model system, as larvae become infected only when feeding on a competent host during their first bloodmeal. For tick-borne diseases, competence has been best studied for *Borrelia burgdorferi* sensu lato (*Bb*sl), which causes Lyme borreliosis. Major reservoirs include several small mammal species, but birds may play an underrecognized role in human risk given their ability to disperse infected ticks across large spatial scales. Here, we provide a global synthesis of the ecological and evolutionary factors that determine the ability of bird species to infect larval ticks with *Bb*sl.

**Location:** Global

**Time period:** 1983 to 2019

**Major taxa studied:** Birds

**Methods:** We compiled a dataset of *Bb*sl competence across 183 bird species and applied meta-analysis, phylogenetic factorization, and boosted regression trees to describe spatial and temporal patterns in competence, characterize its phylogenetic distribution across birds, reconstruct its evolution, and evaluate the trait profiles associated with competent avian species.

**Results:** Half of sampled bird species show evidence of competence for *Bb*sl. Competence displays moderate phylogenetic signal, has evolved multiple times across bird species, and is pronounced in the genus *Turdus*. Trait-based analyses distinguished competent birds with 80% accuracy and show that such species have low baseline corticosterone, exist on both ends of the pace-of-life continuum, breed and winter at high latitudes, and have broad migratory movements into their breeding range. We use these trait profiles to predict various likely but unsampled competent species, including novel concentrations of avian reservoirs within the Neotropics.

**Main conclusion:** Our results can generate new hypotheses for how birds contribute to the dynamics of tick-borne pathogens and help prioritize surveillance of likely but unsampled competent birds. Our findings further emphasize that birds display underrecognized variation in their contributions to enzootic cycles of *Bb*sl and the broader need to better consider competence in ecological and predictive studies of multi-host pathogens.

## Introduction

As most emerging infectious diseases originate in animals (Jones *et al*., 2008), diverse efforts have aimed to predict reservoir hosts and arthropod vectors of zoonotic pathogens (Morse *et al*., 2012; Babayan *et al*., 2018). Because such predictions can guide surveillance and interventions, they are necessary steps toward a preemptive approach to minimizing pathogen spillover risks (Han & Drake, 2016; Becker *et al*., 2019c). For example, PCR and serological data on Nipah virus from bats were leveraged to prioritize field sampling targets across India in response to a novel human outbreak (Plowright *et al*., 2019a), and similar approaches have been applied to identify hosts of filoviruses, rodent zoonoses, and Zika virus (Han *et al*., 2015, 2016, 2019). However, PCR and serological data better reflect host exposure rather than competence, the ability of a host to transmit a pathogen to a new host or vector (Gervasi *et al*., 2015). Greater attention to competence could thus improve reservoir host predictions (Becker *et al*., 2020b).

Host competence is an individual-level and continuous trait encompassing infection processes that occur within the host following exposure: susceptibility to infection, pathogen development, and pathogen survival until transmission (Merrill & Johnson, 2020). This individual heterogeneity mediates intra- and inter-specific variation in competence, which can produce species with disproportionate contributions to pathogen transmission (VanderWaal & Ezenwa, 2016). For example, American robins (*Turdus migratorius*) are on average more competent for West Nile virus than other bird species, infecting up to 71% of mosquito vectors despite low relative abundance in avian communities (Kilpatrick *et al*., 2006). Similarly, transmission of many helminths is dominated by single host species among small mammals (Streicker *et al*., 2013). Identifying the broader ecological or evolutionary drivers of competence could identify how different species contribute to pathogen transmission (Downs *et al*., 2019).

Competence can be difficult to quantify, as infection status of the donor and recipient host must be known or directional transmission events must be inferred (Archie *et al*., 2009; Martin *et al*., 2016). However, many tick-borne diseases facilitate quantifying competence, as tick larvae hatch free of some pathogens and only become infected with their first bloodmeal (Richter *et al*., 2011). Fed larvae are often collected from wild hosts and tested for infection to establish host-to-vector transmission; however, such data can only approximate competence. Instead, xenodiagnostic experiments provide ideal evidence, as uninfected larvae are fed on infected hosts, often allowed to molt (assuring transstadial transmission), and then tested for pathogen presence to infer competence (i.e., proportion of infected ticks) (Brunner *et al*., 2008).

In this context, inter-specific variation in competence has been best studied for *Borrelia burgdorferi* sensu lato (*Bb*sl), which causes Lyme borreliosis. *Bb*sl is transmitted to humans by nymphal and adult *Ixodes* ticks (Hofhuis *et al*., 2017; Eisen, 2020) and has infection foci across the northern hemisphere and parts of Latin America (Kurtenbach *et al*., 2006; Ivanova *et al*., 2014). Lyme borreliosis is the most common vector-borne disease in the United States (i.e., Lyme disease; (Schwartz, 2017), where fast-lived rodent species, specifically white-footed mice (*Peromyscus leucopus*) and eastern chipmunks (*Tamias striatus*), infect high proportions of larvae and are the most competent mammals (LoGiudice *et al*., 2003; Huang *et al*., 2013; Ostfeld *et al*., 2014). By infecting most naïve vectors, these species contribute disproportionally to the production of infectious nymphs and to human risk (Mather *et al*., 1996; Ostfeld *et al*., 2006).

In contrast to mammals, birds likely play an underrecognized role in the global ecology of *Bb*sl, given that the capacity for flight and long-distance migration can allow avian hosts to disperse infected ticks across continents (Smith Jr *et al*., 1996; Ishiguro *et al*., 2005; Dubska *et al*., 2009; Norte *et al*., 2020). Migratory birds transport 50–175 million ticks across Canada each spring (Ogden *et al*., 2008), and the physiological stress of migration itself may help drive reactivation of latent *Bb*sl infection (Gylfe *et al*., 2000). Individual studies have suggested some birds have important contributions to enzootic maintenance of *Bb*sl (Brinkerhoff *et al*., 2011; Mysterud *et al*., 2019). For example, birds dominate transmission of *B. garinii* and *B. valaisiana* to larvae across Europe (Hanincová *et al*., 2003; Comstedt *et al*., 2006) and contribute to enzootic cycles of the primarily rodent genospecies, *B. afzelli* (Franke *et al*., 2010). In North America, *Bb* sensu stricto infects both rodents and birds; however, as with mammals, birds seemingly display interspecific variation in competence. For example, American robins infect up to 90% of naïve larvae (Richter *et al*., 2000), whereas gray catbirds (*Dumetella carolinensis*) and veeries (*Catharus fuscescens*) infect fewer larvae and thus have lower competence (Anderson *et al*., 1986; Mather *et al*., 1989; Ginsberg *et al*., 2005). High tick burdens of some bird species, such as ground foragers, may allow birds to contribute more to *Bb*sl transmission than some rodents (Wright *et al*., 2000; Loss *et al*., 2016). Many birds capable of infecting larvae are also common in suburban and urban habitats (Battaly & Fish, 1993; Hamer *et al*., 2012), which could increase human exposure to infectious nymphs (Mead *et al*., 2018). Yet despite opportunities for birds to play key roles in the global distribution of *Bb*sl and Lyme borreliosis risk, inter-specific drivers of reservoir competence across bird species have not yet been systematically identified.

Given the potential for birds to play important roles in the dynamics of *Bb*sl, we here compile a comprehensive, global dataset on avian competence and assess its ecological and evolutionary drivers. We first describe spatial and temporal patterns in competence, characterize its phylogenetic distribution across birds, and reconstruct its evolution. We then use a flexible machine learning algorithm to evaluate the trait profiles of competent avian species and predict unsampled reservoirs. For the latter, such data science approaches circumvent many issues associated with traditional hypothesis testing (e.g., a large number of predictors, complex interactions, non-randomly missing covariates) and can uncover new and surprising patterns in data, thereby developing testable hypotheses (Hochachka *et al*., 2007). Our work therefore aimed to identify the ecological and evolutionary drivers of avian competence while also generating predictions of likely novel *Bb*sl reservoirs and directions for future studies of tick-borne disease.

## Methods

### Competence data

To collate data on avian competence for *Bb*sl, we searched Web of Science, PubMed, and CAB Abstracts with the following string: (“reservoir competence” OR “host competence” OR prevalence) AND (bird* OR Aves) AND (larva* OR tick* OR arthropod*) AND (“Lyme disease” OR *Borrelia* OR “*B. burgdorferi*”). Using a systematic protocol (Fig. S1), we only included xenodiagnostic experiments and field studies that tested engorged larvae for *Bb*sl. We caution that the latter ignores both transstadial transmission and infection in the host. Although transstadial transmission is well established for *Bb*sl in *Ixodes* ticks (Burgdorfer & Gage, 1986), the absence of *Bb*sl in engorged larvae could simply result from lack of infection in the wild host rather than poor competence (Brunner *et al*., 2008). Field-based measures thus only approximate competence, and failure to detect *Bb*sl in fed larvae requires either testing hosts or experimental validation; however, species to target for both approaches can be prioritized by the predictive methods employed here. From our systematic search, we excluded studies that only tested larvae for non-*Bb*sl *Borrelia* (e.g., *B. lonestari*), only tested nymph or adult ticks, pooled ticks by life stage, only tested wild birds themselves (e.g., blood), or pooled competence across bird species.

We identified 102 studies for inclusion, from which we recorded the sampling country and coordinates (or used centroids of reported regions), sampling months and years, bird and tick species, number of sampled and *Bb*sl-positive larvae, *Bb*sl genospecies, and assessment type (experimental trial or testing attached larvae). These studies encompassed 183 bird species for which engorged larval ticks have been tested for *Bb*sl. Each record (*n*=1069) was a test of a bird– tick–*Bb*sl association over space and time, and most studies contributed multiple lines of data (88/102). Most data were from *Ixodes* ticks (90.11%, mostly *I. ricinus* and *I. scapularis*), with the remainder unstated (1.24%) or from *Haemaphysalis* (4.85%), *Hyalomma* (3.71%), or *Amblyomma* (0.10%). Whereas tick genera other than *Ixodes* are unlikely *Bb*sl vectors (Breuner *et al*., 2020), we retained these records as they still indicate transmission from competent birds.

### Meta-analysis of larval infection prevalence

We used a phylogenetic meta-analysis to quantify heterogeneity in competence, the prevalence of *Bb*sl in bird-fed larvae (*n*=964, Fig. S2; some studies only reported presence of infected larvae). We obtained a phylogeny of our 183 bird species from the Open Tree of Life with the *rotl* package in R and used the *ape* package to calculate branch lengths (Paradis *et al*., 2004; Michonneau *et al*., 2016). We then used the *metafor* package to estimate logit-transformed proportions and sampling variances (Viechtbauer, 2010). We used restricted maximum likelihood to fit a random-effects model (REM), which included a species-level random effect (the covariance structure used the phylogenetic correlation matrix), observation nested in a study-level random effect, and weighting by sampling variances to account for sample size. Estimates of variance components were used to derive *I^2^*, the contribution of true heterogeneity to total variance in competence, and to partition variance attributed to each random effect. For avian species, we also calculated phylogenetic heritability (*H^2^*) (Nakagawa & Santos, 2012).

To assess spatial and temporal variation in competence, we fit mixed-effects models (MEMs) with the same random effects to the data describing *Bb*sl prevalence in engorged larvae from only wild birds (*n*=922). Covariates included geographic region (*n*=922), latitude (*n*=778), year (*n*=917), and season (*n*=654). As many studies pooled data over time, we used the sampling year or the mid-point sampling year. We coded season as binary covariates (winter, spring, summer, fall; *n*=10 records were from the southern hemisphere), as studies often reported data per season or pooled across seasons. Given the differences in sample size between predictors, we used Akaike information criterion (AICc) to compare two sets of MEMs (Burnham & Anderson, 2002): (*i*) space and year and (*ii*) space and season. Comparisons included an intercept-only model, and we derived a pseudo-*R^2^* as the proportional reduction in the summed variance components per MEM compared with that of an equivalent REM (López-López *et al*., 2014).

### Phylogenetic analyses

We next aggregated tick–*Bb*sl data per avian species to assess phylogenetic patterns in competence as a simplified, binary trait. Using the *caper* package, we calculated the *D* statistic, where 1 indicates a phylogenetically random trait distribution and 0 indicates phylogenetic clustering under a Brownian motion model of evolution (Fritz & Purvis, 2010). Significant departure from either model was quantified using a randomization test. However, because traits such as competence may also arise under a punctuated equilibrium model of evolution, we next used a graph-partitioning algorithm, phylogenetic factorization, to flexibly identify clades with significantly different propensity to be competent at various taxonomic depths. We used the *taxize* package to obtain a taxonomy from the NCBI database (Chamberlain & Szöcs, 2013) and used the *phylofactor* package to partition competence as a Bernoulli-distributed response in a generalized linear model (Washburne *et al*., 2019). To account for variable study effort, we used the *rwos* package to quantify the number of citations per species in Web of Science and used the square-root transformed values as weights (Han *et al*., 2016; Plowright *et al*., 2019a). We determined the number of significant phylogenetic factors (clades) using a Holm’s sequentially rejective 5% cutoff for the family-wise error rate. Lastly, to assess whether phylogenetic patterns in competence could stem from study effort alone, we performed a secondary analysis to partition Web of Science citation counts for each avian species as a quasi-Poisson response.

To investigate the evolution of competence across bird species, we used the *ape* package and maximum likelihood to reconstruct the ancestral character state (Paradis *et al*., 2004). We compared an equal-rate and an all-rates-different model of evolution with AIC (Schluter *et al*., 1997) and used the most competitive model to perform stochastic character mapping with Markov chain Monte Carlo (*n*=1000) using the *phytools* package (Revell, 2012). We displayed mean posterior probabilities of competence across our sampled avian phylogeny.

### Trait-based analyses

We compiled avian traits from EltonTraits (Wilman *et al*., 2014), the Amniote Life History database (Myhrvold *et al*., 2015), International Union for the Conservation of Nature (Baillie *et al*., 2004), and HormoneBase (Vitousek *et al*., 2018). Traits included diet composition and breadth, foraging strata, life history (e.g., maximum lifespan, clutch size, fledging age, clutches per year), morphology (e.g., adult mass, hatching weight), maximum elevation, global population trend, and physiology (i.e., baseline corticosterone; CORT). Using distribution maps from BirdLife International and the Handbook of the Birds of the World (BirdLife International & NatureServe, 2014), we derived total range size, latitude of the centroids of breeding and non-breeding ranges, and mean migration distance (greater circle distance between these centroids). We also quantified migratory dispersion, the extent to which species inhabit larger (positive) or smaller (negative) areas in the non-breeding season relative to breeding range size, and made binary covariates for migratory strategy (resident, full migrant, partial migrant) (Gilroy *et al*., 2016). For resident species, migration distance and dispersion were set to zero. To represent avian taxonomy, we included binary covariates for each family and any clades identified by phylogenetic factorization; we also included a binary covariate for the Passeriformes (166/183 species). We also derived evolutionary isolation with the *picante* package (Kembel *et al*., 2010). We again used Web of Science citation counts per species to approximate study effort. We transformed continuous predictors that spanned orders of magnitude and excluded those with high homogeneity or missing values for over 80% of birds. We compiled features for our 183 sampled birds and additional unsampled avian species to predict likely but undetected competent hosts. We limited these out-of-sample species to only the 39 families included in our dataset (4508 bird species). Feature definitions, transformations, and coverage are provided in Table S1.

To identify trait profiles of competent birds and to predict likely novel *Bb*sl reservoirs, we used boosted regression trees (BRTs) to fit a predictive model relating binary competence to a predictor matrix of avian traits (Elith *et al*., 2008). BRTs were trained to maximize classification accuracy by learning patterns of features that best distinguish competent and non-competent species. BRTs generate recursive binary splits for randomly sampled predictors, and successive trees are built using residuals of the prior best-performing tree as the new response. Boosting generates an ensemble of linked trees, where each achieves increasingly more accurate classification. Prior to analysis, we randomly split our data into training (90%) and test (10%) datasets while preserving the proportion of positive labels. Models were then trained with the *gbm* package (Ridgeway, 2006), with a maximum of 30000 trees, a learning rate of 0.0001, and an interaction depth of three (Elith *et al*., 2008). BRTs used a Bernoulli error distribution and 10-fold cross-validation, and we used the *ROCR* and *hmeasure* packages to quantify three measures of classification accuracy: area under the receiver operator curve (AUC), sensitivity, and specificity (Sing *et al*., 2005). As BRT results can depend on random splits between training and test data, we used five partitions to generate a model ensemble (Evans *et al*., 2017). To diagnose if trait profiles of competent birds are driven by study effort, we ran a secondary BRT ensemble that modeled Web of Science citation counts as a Poisson response (Plowright *et al*., 2019a).

After assessing accuracy of our BRTs against test data, we applied our model ensemble to the full trait dataset of 4691 avian species (183 sampled and 4508 unsampled species) to generate mean probabilities of competence. This allowed us to differentiate predictions that signal false negatives (i.e., sampled species without evidence of competence) and those denoting undetected reservoirs (i.e., unsampled species). To assess phylogenetic signal in predictions, we estimated Pagel’s λ in the logit-transformed probabilities with the *caper* package (Orme, 2013) and applied phylogenetic factorization to identify clades of particularly likely competent birds. Lastly, to guide surveillance of these false negatives and undetected competent reservoirs, we mapped the distributions of species with mean predicted probabilities of over 50% and 60%.

## Results

### Avian competence on a global scale

Half of all sampled bird species were competent for *Bb*sl (91/183). Only nine bird species had experimental evidence of competence (i.e., xenodiagnosis), and all 91 competent species (with the exception of *Gallus gallus*) had larvae from wild birds test positive (Fig. S2). For records reporting *Bb*sl genospecies from bird-fed larvae, our global data were dominated by *B. garinii* (31%), *B. valaisiana* (20%), *Bb*ss (16%), and *B. afzelii* (15%), representing data biases toward Europe (59%) and North America (37%); 3% of data were from Eastern Asia, whereas 1% were from South America (Fig. 1A). However, these 91 competent bird species were broadly distributed across the Americas, Africa, Asia, and Oceania throughout their annual cycles.

**Figure 1.**
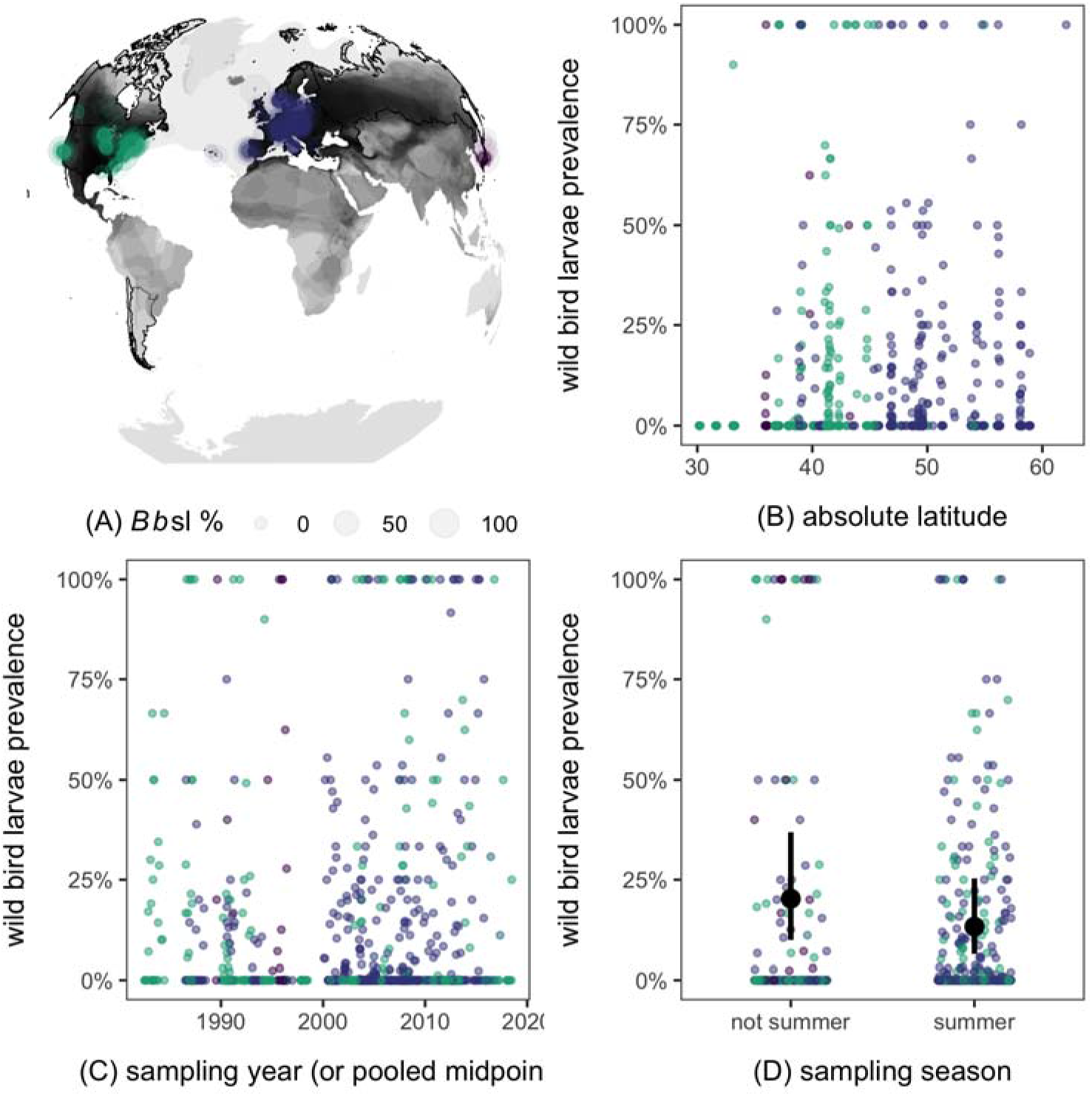
Global distribution of *Bb*sl prevalence in tick larvae sampled from wild birds. (A) Sampled countries are shown in black border with spatiotemporal bird–tick–*Bb*sl prevalences, which are sized by magnitude and colored by georegion. Shapefiles of the 91 competent species are derived from BirdLife International and Handbook of the Birds of the World and are overlaid in black (BirdLife International & NatureServe, 2014). Larval infection prevalence is plotted by absolute latitude (B), year (C), and season (D), with each point representing a bird–tick–*Bb*sl association; points are colored by region and jittered to reduce overlap. Black points and lines (D) display predicted means and 95% confidence intervals from the top MEM (Table S3).

We observed significant heterogeneity in bird-fed larvae *Bb*sl prevalence (*I^2^*=0.76, *Q_963_*=2970, *p*<0.0001). Avian species accounted for more of this variation (*I^2^_species_*=0.31) than study (*I^2^_study_*=0.19) or individual record (*I^2^_observation_*=0.26), resulting in moderate phylogenetic signal (*H^2^*=0.40). Given the stronger effect of avian phylogeny, we found no effect of study-level predictors such as space (Fig. 1B) or year (Fig. 1C) on the proportion of larvae infected by wild birds (Table S2). Seasons were also mostly uninformative (Table S3), but prevalence in bird-infected larvae was weakly lower during summer (Akaike weight=0.44, *R^2^*=0.01; Fig. 1D).

### Evolutionary patterns in competence

We next considered competence as an intrinsic binary trait per species. We estimated intermediate phylogenetic signal in the ability of birds to transmit *Bb*sl to larvae (*D*=0.78), indicating significant phylogenetic clustering between randomness (*p*<0.001) and a Brownian motion model of evolution (*p*<0.001). After controlling for study effort, phylogenetic factorization identified one clade, the genus *Turdus*, as having a significantly greater likelihood of including competent species when compared to other avian taxa (Fig. 2). In particular, all but one sampled member of this clade displayed the ability to infect larval ticks with *Bb*sl (92%).

**Figure 2.**
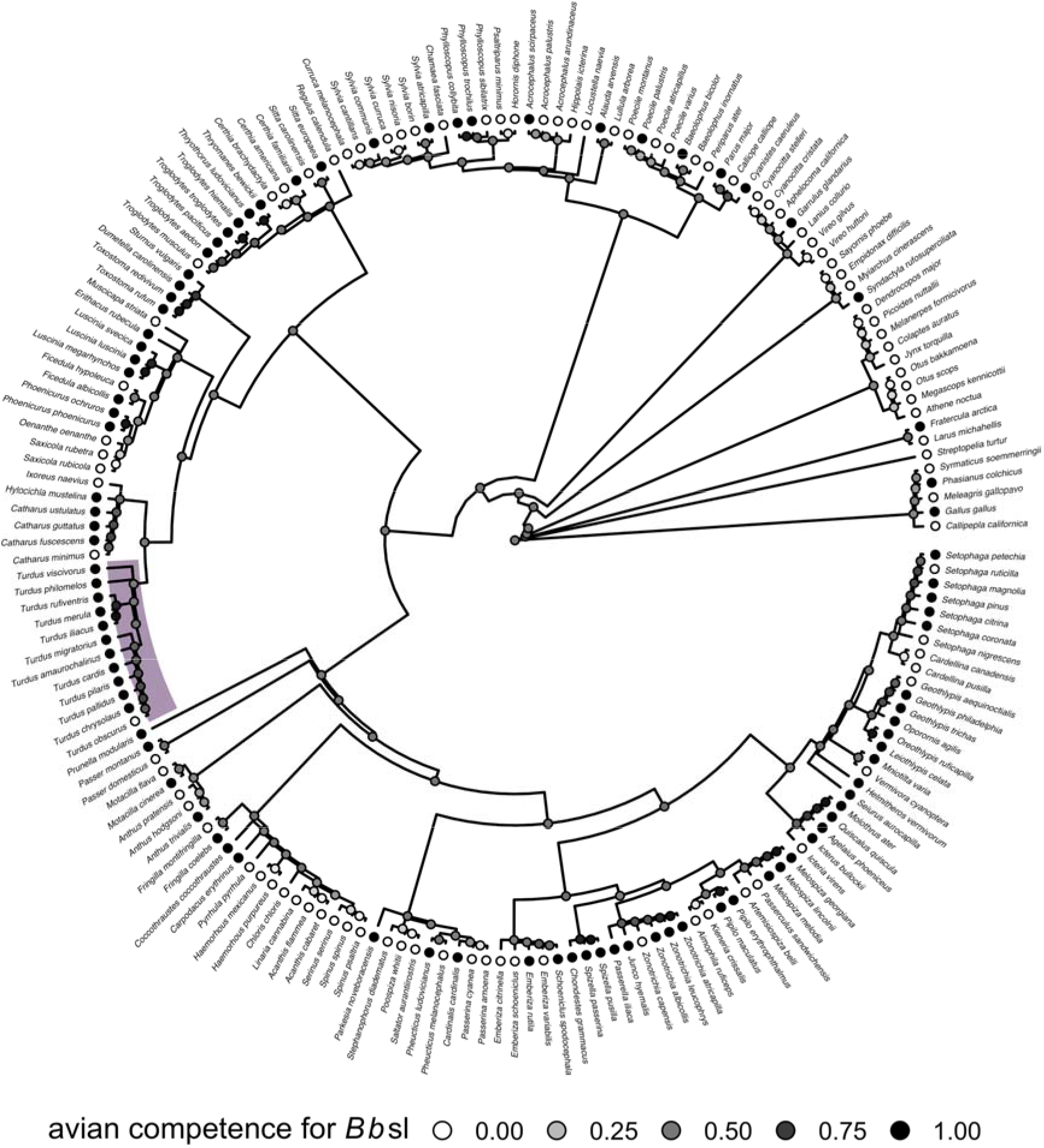
Phylogenetic patterns in avian competence for *Bb*sl. The avian phylogeny displays observed species binary competence, and highlighted clades are those with significantly different competence from the paraphyletic remainder using phylogenetic factorization. Nodes show the mean posterior probabilities of being competent estimated from stochastic character mapping.

Our secondary analysis identified seven taxa as having significantly different Web of Science citation counts (Fig. S3), five of which were heavily studied species: *Gallus gallus* (4139), *Parus major* (3736), *Sturnus vulgaris* (2509), *Passer domesticus* (2162), and *Ficedula hypoleuca* (1500). Disproportionately studied taxa also included the Phasianidae (x□=609) and a subclade of the Icteridae (genera *Quiscalus, Molothrus*, and *Agelaius;* x□=545). Taxonomic patterns in Web of Science citation counts did not overlap with taxonomic patterns in avian competence, suggesting that the latter were not driven by variable study effort (Fig. S3).

The evolution of avian competence was best described by an equal-rate model (*w_i_*=0.73). Stochastic character mapping suggested equal transitions from non-competent to competent and from competent to non-competent (Fig. 2), with the ancestral state being equivocal. Competence was gained within the Turdidae as well as the Mimidae, Passerellidae, and Troglodytidae, with both gains and losses within the Parulidae. We also observed a clear loss of competence within the majority of the Carduelinae as well as within the Corvidae, Picidae, and Strigidae (Fig. 2).

### Trait profiles of competent birds

Our BRT models distinguished competent from non-competent birds with moderate accuracy (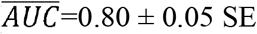; 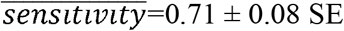; 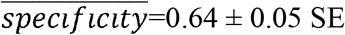; Fig. 3A). Some top features for describing *Bb*sl-competent species included physiology (i.e., baseline CORT), life history (i.e., fledging age, maximum lifespan, birth and fledging weight, egg mass, incubation time, clutch size), migration (dispersion, distance), geography (breeding and non-breeding latitude, maximum elevation, geographic range size), evolutionary isolation, and study effort. BRTs identified our phylofactorization clade (i.e., the *Turdus* genus) alongside the Turdidae family and passerines more generally as the only taxonomic features with non-trivial importance (i.e., <1%), and foraging traits were generally uninformative (Fig. 3B, Table S4).

**Figure 3.**
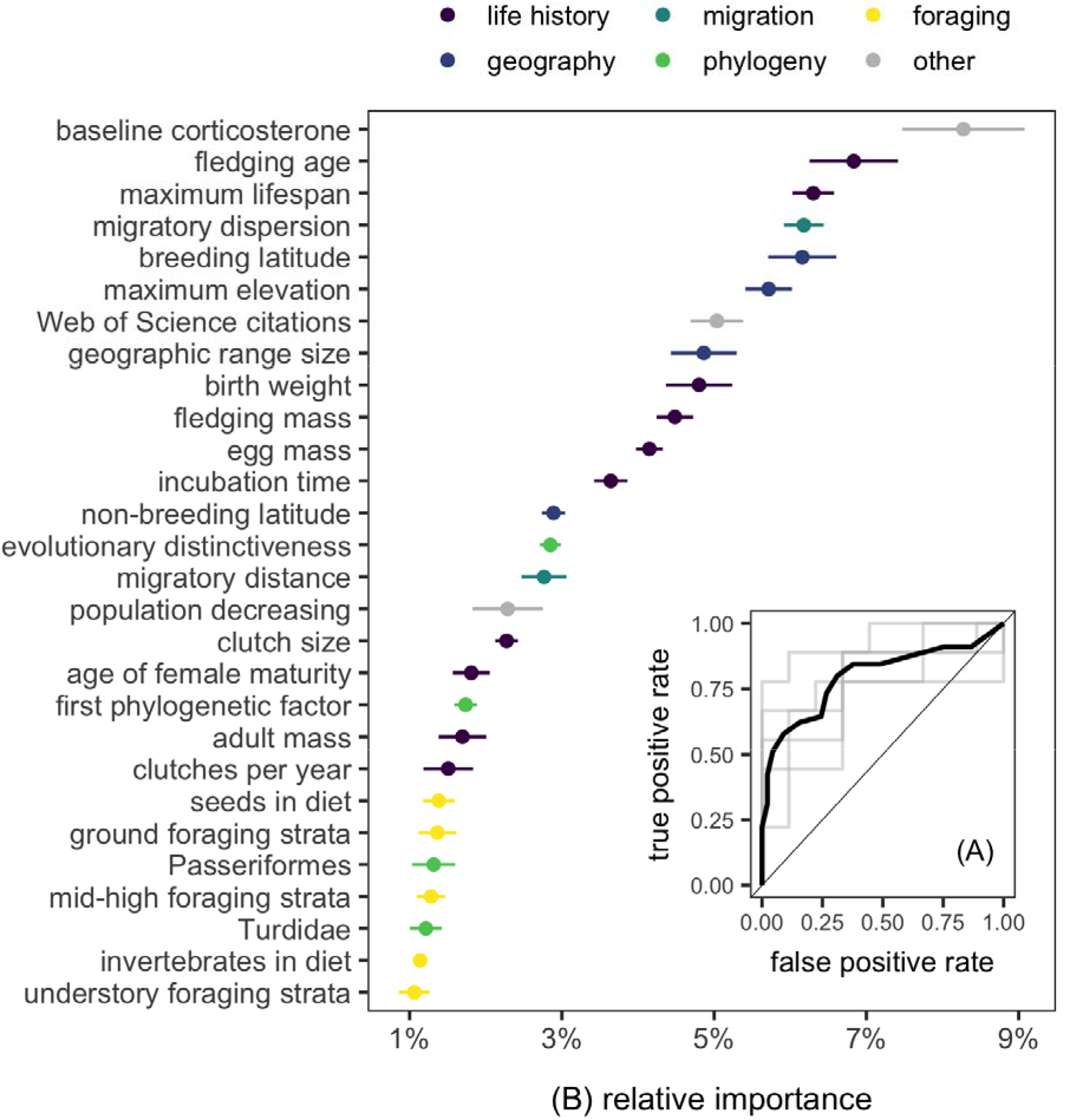
BRT performance in identifying traits predictive of avian competence for *Bb*sl and their relative importance across five random partitions of training and test datasets. (A) Accuracy is shown by the receiver operator curves obtained from 10-fold cross-validation on test data. Grey lines show curves from each partition, whereas the black line displays the mean. (B) Relative importance per feature is shown as the mean and standard error across training and test data partitions. Only features with mean importance greater than 1% are shown (Table S4).

Physiologically, *Bb*sl-competent birds have lower baseline concentrations of CORT, the main avian glucocorticoid (Fig. 4). Competent species were also described by either extreme of the pace-of-life continuum. On the one hand, competent birds have shorter incubation times and young that are smaller and fledge earlier than non-competent counterparts. On the other hand, competent birds also had longer lifespans, smaller clutches, and larger eggs and body size. Geographically, competent birds tend to breed and winter at higher latitudes, have broader distributions, and occupy lower elevations, and their populations are stable or increasing. Species with negative migratory dispersion were more likely to be competent, indicating such hosts have larger breeding ranges and greater diversity of migratory movements from their wintering to breeding grounds. Competence was less strongly related to mean migratory distance, with intermediate-distance migrants being most likely to transmit *Bb*sl to larvae. Members of the genus *Turdus*, the family Turdidae, and the Passeriformes as a whole were all more likely to be competent. Additionally, although foraging traits generally had low relative importance, competent birds were more likely to be granivorous and ground-foraging species. Lastly, well-studied species were also more likely to be competent. However, our secondary BRTs showed that citations were not predictable by these same traits 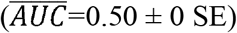, suggesting that the trait profile of a competent bird is not confounded by the ecological traits of well-studied hosts.

**Figure 4.**
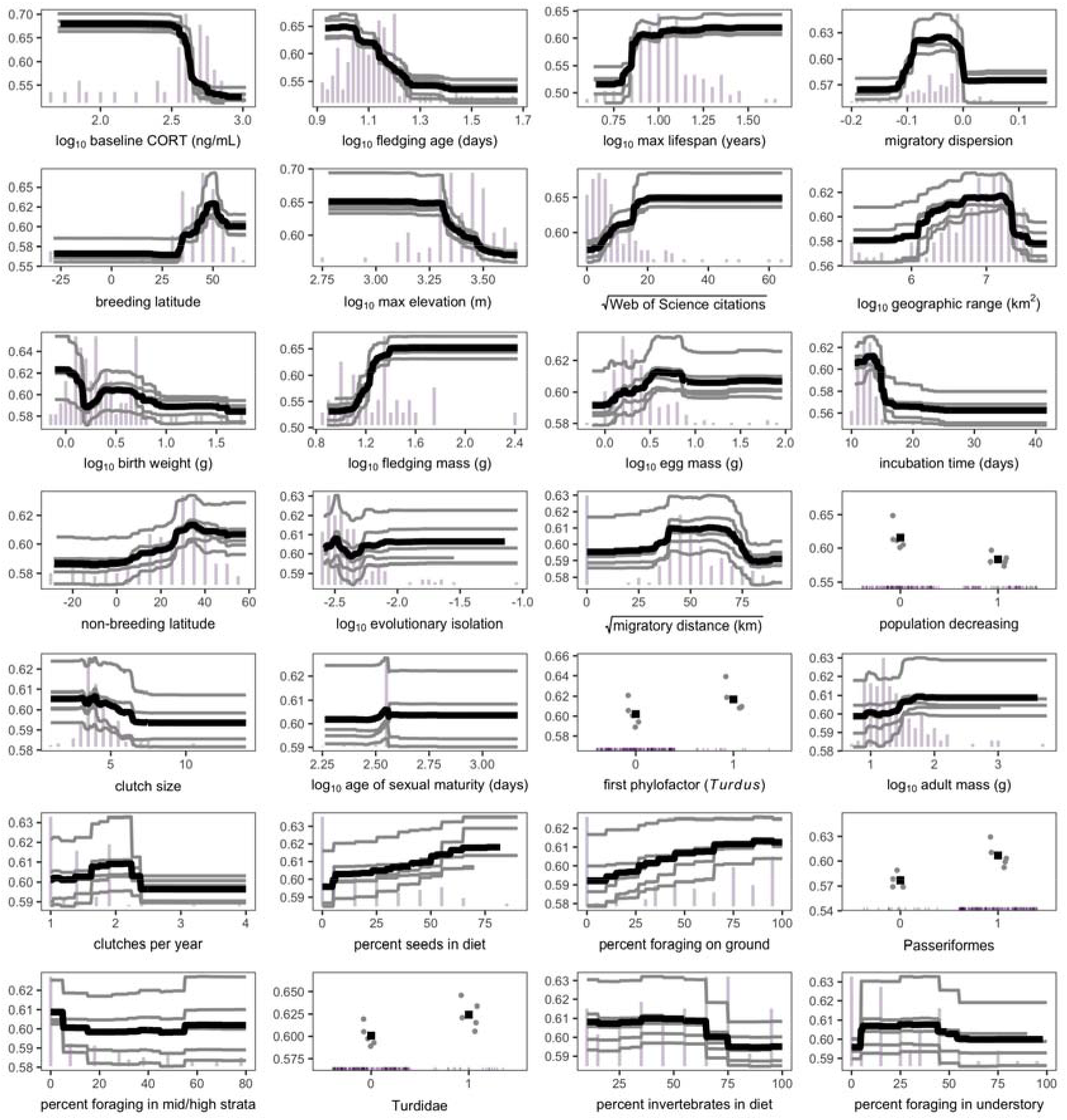
Trait profile of a *Bb*sl-competent avian species. Partial dependence plots of the top predictors across BRTs applied to five random partitions of training and test data are shown ordered by relative importance (>1%). Grey lines or point show the marginal effect of a given variable for predicting *Bb*sl competence from each random data partition, whereas the black lines or squares display the average marginal effect. Histograms and rug plots display the distribution of continuous and categorical predictor variables, respectively, across the 183 sampled birds.

Applying our BRT ensemble to trait and taxonomic data across the 39 sampled avian families revealed at least 21 undiscovered species that could be prioritized for *Bb*sl surveillance based on feature similarity to known competent reservoirs (Fig. 5A). We observed strong phylogenetic signal in mean predicted probabilities (λ=0.94), indicating the influential traits revealed by our BRTs are likely driven by clades with high potential for competence (Fig. 5B). Phylogenetic factorization found nine such clades with distinct model predictions (Table S5).

**Figure 5.**
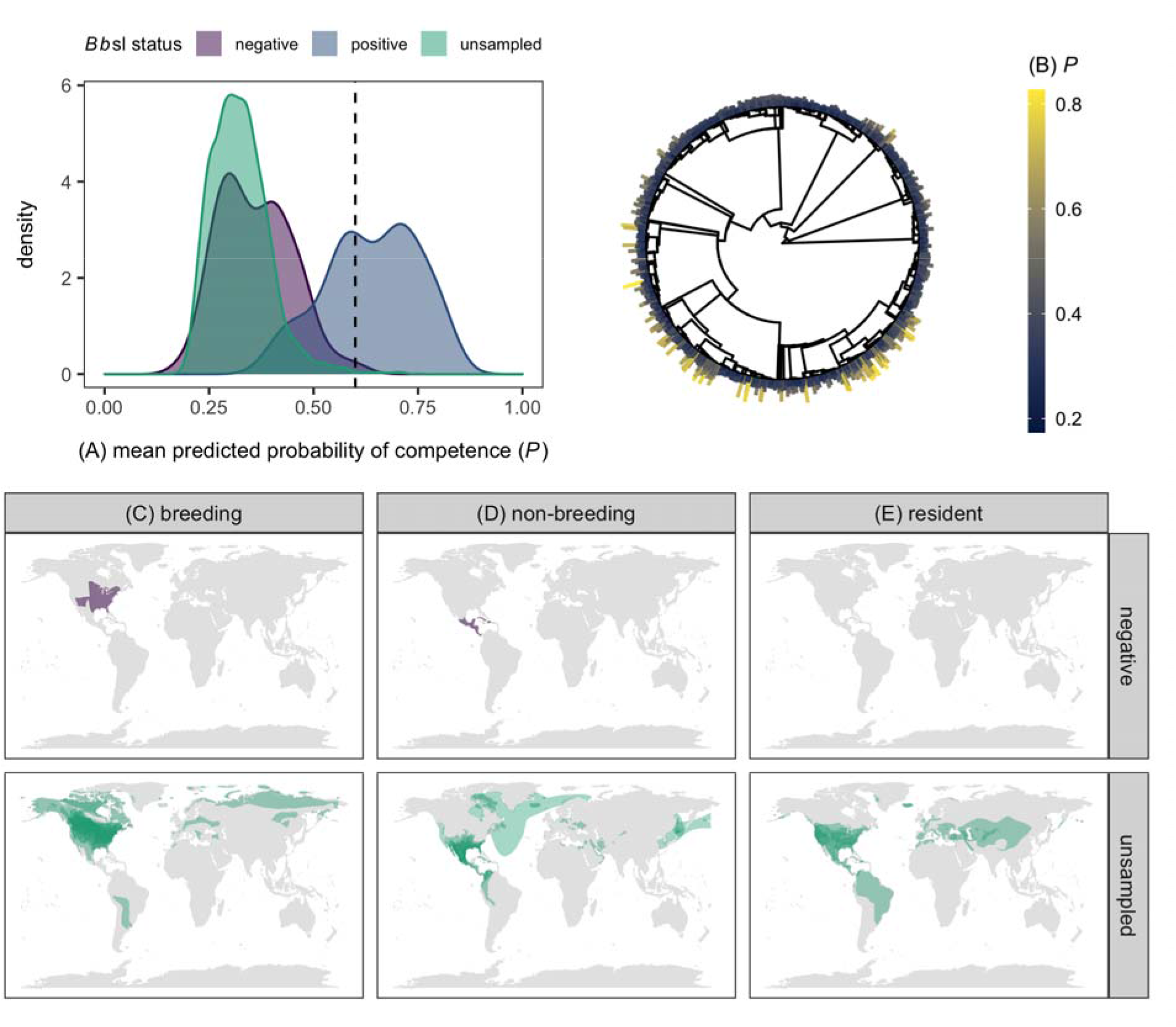
Distribution of the mean predicted probabilities of *Bb*sl competence across the 39 sampled avian families. (A) Density plots show predictions for currently negative, positive, and unsampled species, and these propensity scores are also shown across the avian phylogeny (B). Distributions of species within a mean probability of over 60% are shown by their breeding (C), non-breeding (D), and resident ranges (E). Shapefiles are from BirdLife International and the Handbook of the Birds of the World (BirdLife International & NatureServe, 2014). Figure S4 displays these geographic distributions with a less conservative prediction cutoff of 50%.

The geography of likely high-probability reservoirs revealed potential hotspots of competent birds across their breeding (Fig. 5C), non-breeding (Fig. 5D), and resident (Fig. 5E) ranges. Our BRTs suggested at least one likely false negative, the Indigo bunting (*Passerina cyanea*; Sonenshine *et al*., 1995; Kinsey *et al*., 2000; Schneider *et al*., 2015). Likely but unsampled competent reservoirs included the American goldfinch (*Spinus Iristis*), Harris’s sparrow (*Zonotrichia querula*), Abert’s towhee (*Melozone aberti*), yellow-headed blackbird (*Xanthocephalus xanthocephalus*), western meadowlark (*Sturnella neglecta*), northern mockingbird (*Mimus polyglottos*), Brewer’s blackbird (*Euphagus cyanocephalus*), and vesper sparrow (*Pooecetes gramineus*) in North America; Townsend’s warbler (*Setophaga townsendi*), scarlet tanager (*Piranga olivacea*), eastern bluebird (*Sialia sialis*), Louisiana waterthrush (*Parkesia motacilla*), red-eyed vireo (*Vireo olivaceus*), grasshopper sparrow (*Ammodramus savannarum*), Acadian flycatcher (*Empidonax virescens*), and clay-colored thrush (*Turdus grayi*) across the Americas; the corn bunting (*Emberiza calandra*) across Eurasia; the horned lark (*Eremophila alpestris*) across the northern hemisphere; and the thick-billed murre (*Uria lomvia*) and rhinoceros auklet (*Cerorhinca monocerata*) across pelagic zones. Species with probabilities above 50% (96^th^ percentile of predictions) are included in Table S6 and shown in Figure S4; this cutoff found another three likely false negatives and 74 likely but unsampled competent species.

## Discussion

Competence is increasingly recognized to govern infectious disease dynamics, especially in multi-host communities (Ostfeld *et al*., 2014; Gervasi *et al*., 2015). Efforts to predict new reservoirs of zoonotic pathogens would be improved by identifying the ecological or evolutionary drivers of this trait (Becker *et al*., 2020b). We here applied such an approach to Lyme borreliosis, a model system to study competence for a tick-borne disease. Whereas inter-specific variation in competence has been characterized for mammals (LoGiudice *et al*., 2003; Huang *et al*., 2013; Ostfeld *et al*., 2014), this trait has been understudied across birds, despite their ability to disperse infected ticks across large spatial scales and mounting evidence of competence across avian species (Richter *et al*., 2000; Ginsberg *et al*., 2005; Norte *et al*., 2013). We here demonstrate that *Bb*sl competence can be predicted by the ecological and evolutionary characteristics of birds. Our phylogenetic analyses show that competence has evolved multiple times and is pronounced in the genus *Turdus*. Trait-based analyses distinguished competent avian hosts with 80% accuracy and emphasized that such species have low baseline CORT, occur on either extreme of the pace-of-life continuum, breed and winter at high latitudes and low elevations, and have diverse migratory movements into their breeding range. These patterns can be used to generate testable hypotheses for future studies, and predictions using these trait profiles can help prioritize surveillance of false negatives and likely but unsampled competent avian species. More broadly, these results emphasize birds display underrecognized intra- and interspecific variation in their contributions to the enzootic cycles of this zoonotic pathogen.

Although pathogen transmission inherently occurs from individuals, variation in competence can arise across broader biological scales (Gervasi *et al*., 2015; VanderWaal & Ezenwa, 2016). Our meta-analysis identified high heterogeneity in *Bb*sl prevalence from bird-fed larvae that was better explained by within-study heterogeneity and bird phylogeny than study-level variation and broad spatial or temporal covariates. Greater within-than between-study heterogeneity suggests a greater role for fine-scale environmental covariates, such as local densities of ticks or alternative hosts, in shaping vector feeding behavior and infection prevalence in avian hosts that in turn affect prevalence in bird-fed larvae (Kilpatrick *et al*., 2017). Additionally, moderate contributions of avian phylogeny in our meta-analysis was mirrored by intermediate phylogenetic signal in our species-level analysis. Together, these analyses suggest that while intraspecific variation in host competence occurs over space and time (Norte *et al*., 2020), the ability to transmit *Bb*sl can also be considered an intrinsic trait of avian species.

Moderate phylogenetic signal in competence was reflected in numerous gains and losses of this trait across the avian phylogeny, suggesting competence has evolved multiple times to form clades of highly competent birds. Phylogenetic factorization identified one of these clades, the genus *Turdus*, as being especially competent, even after accounting for variable study effort. Our BRTs also identified this genus, and the family Turdidae, as having high likelihood of competence. Of the 12 sampled *Turdus* species, all but one (*T. obscurus*) had fed larvae test positive for *Bb*sl (Ishiguro *et al*., 2000), and two (*T. migratorius* and *T. merula*) have experimentally infected larvae (Richter *et al*., 2000; Norte *et al*., 2013). Given the larger *Turdus* and Turdidae clades, these results suggest unsampled thrushes could also be competent *Bb*sl reservoirs. Thrushes are globally distributed and display diverse migratory strategies, with resident, migratory, and partial migratory species. However, most European *Turdus* are complete or partial migrants (Ashmole, 1962), and photoperiod manipulations suggest migration of at least one species (*T. iliacus*) can reactivate latent *Bb*sl infection (Gylfe *et al*., 2000). Comparative studies across thrushes that vary in geography and migratory strategy could elucidate the generality of this pattern and the broader inter-specific drivers of competence in this taxon.

Our BRTs identified CORT as the top predictor of competence, for which competent birds were more likely to have lower baseline concentrations. Although persistently elevated CORT can be immunosuppressive and amplify competence (Gervasi *et al*., 2017), baseline concentrations have mostly metabolic functions that allow animals to meet energetic demands and respond to adverse conditions (Sapolsky *et al*., 2000). Low baseline CORT could be linked to competence by its association with breeding latitude. Birds tend to have lower baseline CORT at high latitudes, which could facilitate continued breeding in suboptimal habitats (Wingfield & Sapolsky, 2003). This is compatible with our finding that competent species also generally breed at high latitudes. Birds breeding further from the equator show stronger tradeoffs between arms of the immune system, such that high-latitude hosts mount weaker adaptive responses (Ardia, 2007). Because robust *Bb*sl-specific antibody titers limit transmission to naïve ticks (Kurtenbach *et al*., 1994), birds breeding at high latitudes could display weaker antibody defenses that increase competence. Sampling competent birds across latitudinal gradients could characterize such immunity tradeoffs and test if these restrict bird–tick transmission (Becker *et al*., 2019a).

Our trait-based analyses also suggested that competent birds occur on either extreme of the pace-of-life continuum (Stearns, 1983). This possibly contrasts with work on mammals, where fast-lived species (i.e., rapid development and high fecundity at the expense of longevity) are more competent than their slow-lived counterparts (LoGiudice *et al*., 2003; Huang *et al*., 2013; Ostfeld *et al*., 2014). Although many of the top traits predicting competence in birds also reflect fast pace-of-life (e.g., short incubation times, young that are small and fledge early), competent birds were also characterized by long lifespans, large eggs, and small clutches more consistent with a slow life history. This pattern could arise from two competing signals in the data, such that both particularly fast- (e.g., many passerines) and slow-lived species (e.g., the Alcidae and Phasianidae) display evidence of competence. Future tests of pace-of-life variation and competence within orders such as the Passeriformes or Charadriiformes could minimize confounding effects of taxonomy and assess if such patterns have an immunological basis, as suggested for mammalian competence (Previtali *et al*., 2012; Albery & Becker, 2020).

Our BRTs identified several other important predictors related to bird geography and annual cycles, including low elevation, large distributions, and negative migratory dispersion. Rather than indicating physiological processes that facilitate bird–tick transmission, greater likelihood of competence in species at low elevations and with large geographic ranges may indicate greater exposure to questing nymphs that would cause infection in birds. Similarly, positive associations between competence and both breeding and wintering latitude could stem from optimal overlap with tick species (Hahn *et al*., 2016; Hvidsten *et al*., 2020). Although foraging traits were largely uninformative, positive associations between ground foraging and competence likely also reflect greater tick exposure (Loss *et al*., 2016). However, migratory traits could better reflect within-host processes of competence itself. Negative migratory dispersion indicates birds with more diverse movements from their wintering to breeding grounds (Gilroy *et al*., 2016). These more diverse migrations demand large energy expenditures that can impair immunity (Owen & Moore, 2008) and cause latent *Bb*sl infections to reactivate (Gylfe *et al*., 2000). This mechanism could facilitate competent birds arriving at their breeding grounds primed to infect larval ticks. The generally positive association between migratory distance and competence also supports the idea that longer biannual migration, in being more costly, could promote relapse. Future work could test this hypothesis by sampling competent birds across their annual cycles (Marra *et al*., 2015) and linking such data with mathematical models to understand when migratory relapse most increases risk (Becker *et al*., 2020a).

Our analyses also inform surveillance of specific bird species for their contribution to enzootic cycles of *Bb*sl and other tick-borne pathogens. Phylogenetic factorization and our BRTs suggested unsampled *Turdus* thrushes are especially likely to be competent. Some thrushes such as *T. grayi* are known to be parasitized by ticks (Miller *et al*., 2016), whereas others such as *T. torquatus* and *T. nigriceps* have had engorged nymphs test positive for *Bb*sl (Hasle *et al*., 2011; Saracho Bottero *et al*., 2017). Our BRT predictions displayed high phylogenetic signal, identifying clades of especially competent birds, such as the genus *Zonotrichia* as well as the families Alcidae, Mimidae, and Parulidae (Table S5). Some unsampled species in these clades have had blood or nymphs test positive for *Bb*sl, such as *Parkesia motacilla* (Anderson & Magnarelli, 1984). We suggest members of these clades be prioritized for spatiotemporal sampling to identify when and where they are likely to infect ticks (Plowright *et al*., 2019b).

To test these model predictions, we encourage more definitive assessments of competence. Because most of our data included approximations of competence from *Bb*sl in engorged larvae on wild birds, xenodiagnostic experiments could be prioritized for unsampled avian species with high probabilities of being competent to establish bird–tick transmission (Ginsberg *et al*., 2005; Norte *et al*., 2013). As an alternative approach, field surveys could instead assess *Bb*sl infection in not only engorged larvae but also hosts themselves to test whether absence of the pathogen in ticks is due to poor competence or an uninfected host (Newman *et al*., 2015). These increasing data on host competence would also facilitate future analytic efforts. Because *Bb*sl includes genospecies that vary in host specificity (e.g., *Bb*ss infects both rodents and birds, whereas *B. garinii* and *B. valaisiana* are more specialized on the latter), better considering such coevolutionary relationships could improve model performance (Kurtenbach *et al*., 2006; O’Keeffe *et al*., 2020). We pooled *Bb*sl across genospecies due to the relatively small sample of bird species, but models applied to taxonomic subsets of data may generate distinct predictions by reducing noise from hosts infected with other *Bb*sl genospecies.

Lastly, environmental change could play an important role in shaping how known and likely competent birds contribute to *Bb*sl dynamics. Breeding ranges of many birds are shifting north with climate change (Hitch & Leberg, 2007), which could synchronize bird and tick phenologies (Ostfeld & Brunner, 2015). Alternatively, warmer temperatures could facilitate residency, as observed for competent birds such as *Turdus merula* (Vliet *et al*., 2009). If such species become resident where ticks are abundant, sedentary behavior could increase vector exposure and amplify bird–tick transmission. Similarly, several *Bb*sl-competent birds (e.g., *Sylvia atricapilla, Junco hyemalis*) and unsampled but likely reservoirs (e.g., *Spinus tristis*) have shortened migration or become resident in cities (Yeh & Price, 2004; Plummer *et al*., 2015; Bonnet-Lebrun *et al*., 2020). This urban residency could increase or decrease competence depending on factors such as food availability and artificial light at night (Becker *et al*., 2019b; Kernbach *et al*., 2019). Combining sampling of known or likely competent birds across urban-rural gradients or their historic and recent range with mathematical models could forecast how environmental change will alter bird distributions, competence, and contribution to *Bb*sl risk.

In conclusion, we demonstrate that host ability to transmit pathogens to new hosts or vectors can be predicted by the ecological and evolutionary characteristics of bird species in the Lyme borreliosis system. By combing flexible phylogenetic and trait-based analyses, our work generates testable hypotheses for future comparative and theoretical studies of tick-borne disease alongside predictions that can inform bird surveillance efforts for not only *Bb*sl but also similar pathogens (e.g., *Anaplasma, Ehrlichia*) in the context of environmental change. Moving forward, greater attention to the factors that shape competence within and between species could improve our ability to predict and manage reservoir hosts for zoonotic pathogens more broadly.

## Data availability

Data and R code to reproduce the primary analyses are available in Dryad Digital Depository (Becker & Han, 2020): https://datadryad.org/stash/share/4shvYd0_lCUkjEFhO5g1yc95sSLUfq3zbrd79_1J-0Q.

## Competing interests

We declare no competing interests.

## Acknowledgments

We thank BirdLife International for providing avian distribution data, Tao Huang for assistance with data processing, and JP Schmidt for technical advice on BRTs. We also thank Ellen Ketterson, members of the Ketterson lab, and two anonymous reviewers for feedback on this manuscript. DJB was supported by an appointment to the Intelligence Community Postdoctoral Research Fellowship Program, administered by Oak Ridge Institute for Science and Education through an interagency agreement between the U.S. Department of Energy and the Office of the Director of National Intelligence. BAH was supported by the National Science Foundation Ecology and Evolution of Infectious Diseases program (DEB-1717282 and DEB-1619072).

## Supporting information

Additional supporting information may be found online in the Supporting Information section.

### Biosketch

Daniel Becker studies the ecology and evolution of infectious disease, with particular interests in how anthropogenic factors affect infection dynamics in reservoir hosts and how competence data can improve predicting reservoirs of zoonotic pathogens. Barbara Han is a disease ecologist at the Cary Institute of Ecosystem Studies, where she applies machine learning and other modeling in the predictive analytics of zoonotic risk.

## Notes

### Competing Interest Statement

The authors have declared no competing interest.

