## Supplemental Material for "The macroecology and evolution of avian competence for *Borrelia burgdorferi*"

**A. Literature search**
**B. Variation in larval *Bb*sl prevalence**
**C. Trait-based analysis dataset**
**D. Phylogenetic meta-analysis**
**E. Phylogenetic factorization**
**F. Boosted regression trees**

### A. Literature search

Figure S1. PRIMSA diagram documenting the data collection and inclusion process for *Bb*sl competence of avian hosts. We ran all systematic searches in January 2020 and supplemented results by extracting data from references cited in systematically identified studies.

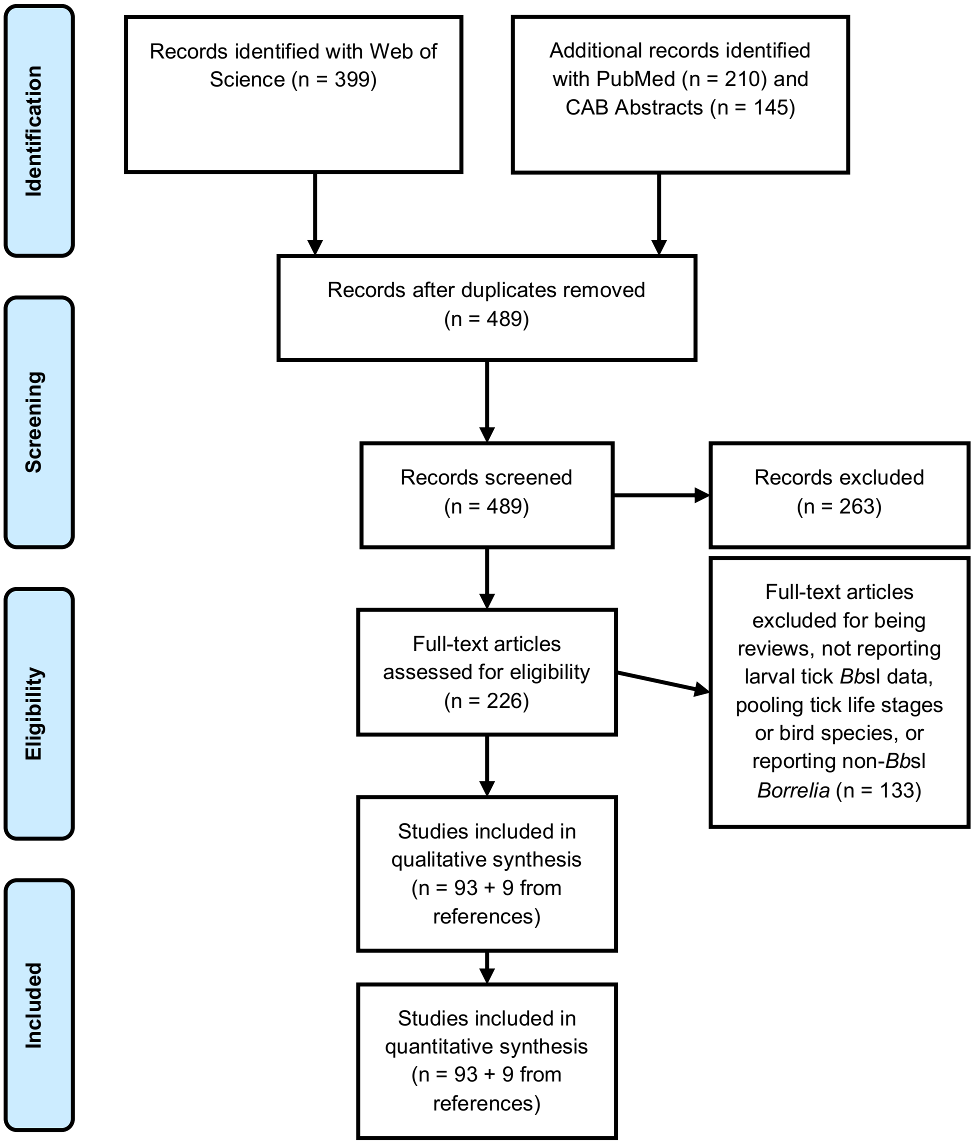

*Included studies*

42. Norte AC, Margos G, Becker NS, Ramos JA, Núncio MS, Fingerle V, et al. Host dispersal shapes the population structure of a tick-borne bacterial pathogen. Molecular Ecology. n/a. doi:10.1111/mec.15336

75. Chvostáč M, Špitalská E, Václav R, Vaculová T, Minichová L, Derdáková M. Seasonal Patterns in the Prevalence and Diversity of Tick-Borne Borrelia burgdorferi Sensu Lato, Anaplasma phagocytophilum and Rickettsia spp. in an Urban Temperate Forest in South

Western Slovakia. International journal of environmental research and public health. 2018;15: 994.

85. Pereira A, Parreira R, Cotão AJ, Nunes M, Vieira ML, Azevedo F, et al. Tick-borne bacteria and protozoa detected in ticks collected from domestic animals and wildlife in central and

southern Portugal. Ticks and Tick-borne Diseases. 2018;9: 225–234. doi:10.1016/j.ttbdis.2017.09.008

**B. Variation in larval *Bb*sl prevalence**

Figure S2. Distribution of *Bb*sl infection prevalence from larval ticks collected from birds. Prevalence estimates per species are sized by the number of larval ticks submitted for testing. Bird species with non-zero prevalence or with experimental evidence of transmitting *Bb*sl to naïve larval ticks were considered to be competent reservoirs (right) in phylogenetic and trait-based analyses. Bird species with consistent zero larval prevalence are displayed on the left. Several birds are listed as competent despite zero larval prevalence (e.g., *Garrulus glandarius*) owing to records where positive larvae were reported without listing total larval sample sizes.

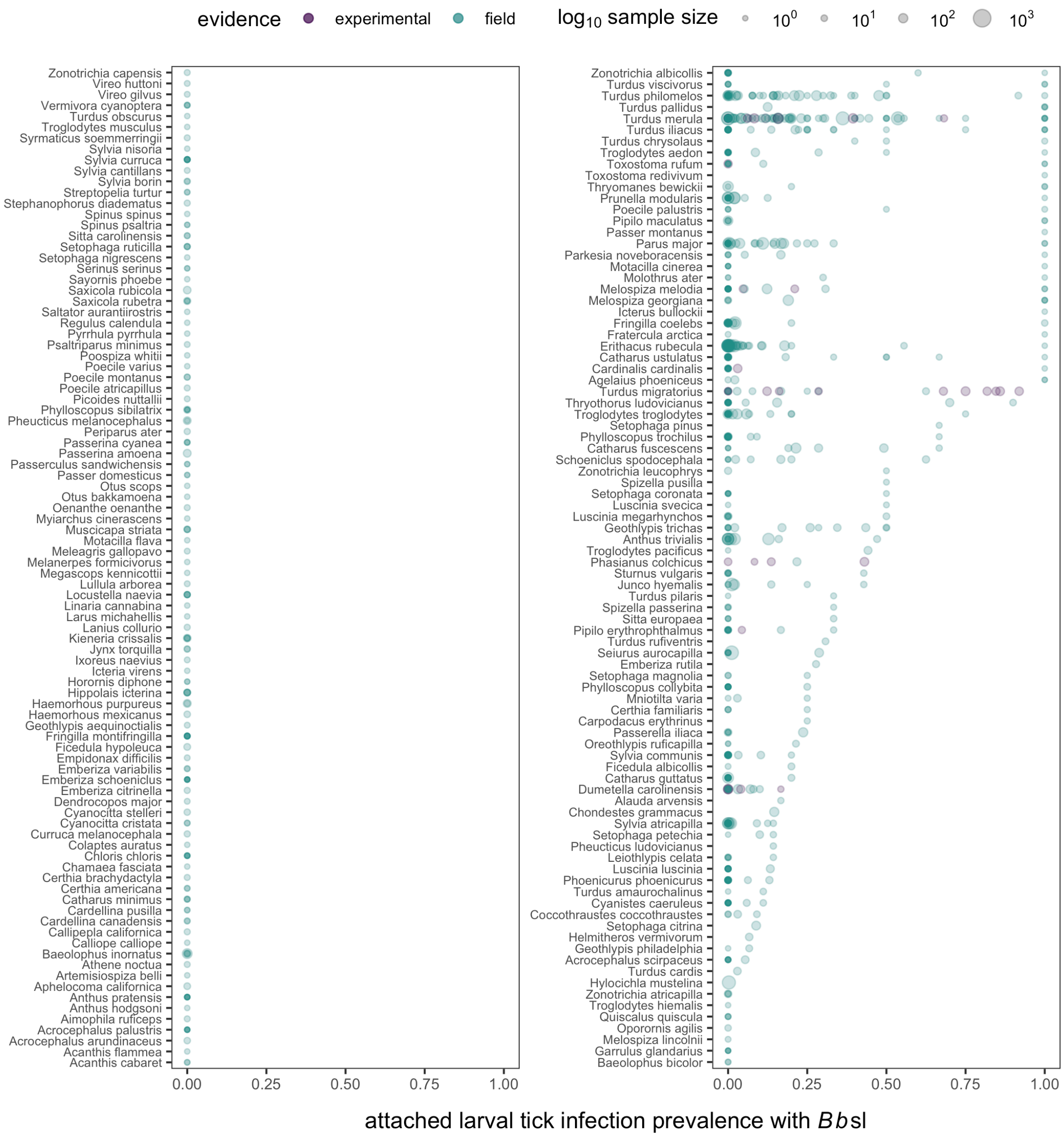

### C. Trait-based analysis dataset

Table S1. Boosted regression tree features, definitions, data type, coverage, any transformations applied, and sources.

| Feature | Definition | Class | Coverage | Transform | Source |
|---|---|---|---|---|---|
| breed_lat | Latitude of breeding distribution centroid | Continuous | 0.98 | NA | [1] |
| diet_diversity | Number of non-zero diet items | Count | 0.46 | NA | [2] |
| Diet.Fruit | % fruit in diet | Proportion | 0.46 | NA | [2] |
| Diet.Inv | % invertebrates in diet | Proportion | 0.46 | NA | [2] |
| Diet.Nect | % nectar in diet | Proportion | 0.46 | NA | [2] |
| Diet.PlantO | % other plants in diet | Proportion | 0.46 | NA | [2] |
| Diet.Scav | % scavenged in diet | Proportion | 0.46 | NA | [2] |
| Diet.Seed | % seeds in diet | Proportion | 0.46 | NA | [2] |
| Diet.Vect | % ectotherms in diet | Proportion | 0.46 | NA | [2] |
| Diet.Vend | % endotherms in diet | Proportion | 0.46 | NA | [2] |
| Diet.Vfish | % fish in diet | Proportion | 0.46 | NA | [2] |
| family_Emberizidae | Species in given family | Binary | 1 | NA | [2,3] |
| family_Fringillidae | Species in given family | Binary | 1 | NA | [2,3] |
| family_Muscicapidae | Species in given family | Binary | 1 | NA | [2,3] |
| family_Paridae | Species in given family | Binary | 1 | NA | [2,3] |
| family_Parulidae | Species in given family | Binary | 1 | NA | [2,3] |
| family_Turdidae | Species in given family | Binary | 1 | NA | [2,3] |
| foraging_diversity | Number of non-zero foraging classes | Count | 0.46 | NA | [2] |
| ForStrat.canopy | % foraging time above canopy | Proportion | 0.46 | NA | [2] |
| ForStrat.ground | % foraging time on ground | Proportion | 0.46 | NA | [2] |
| ForStrat.midhigh | % foraging time in mid to high levels in trees | Proportion | 0.46 | NA | [2] |
| ForStrat.understory | % foraging time below 2m in understory | Proportion | 0.46 | NA | [2] |
| freshwater_system | Does species inhabit a freshwater ecosystem | Binary | 0.98 | NA | [4] |
| incubation_d | Days between laying and hatching | Continuous | 0.84 | NA | [3] |
| IUCN_LC | IUCN Red List Category of Least Concern | Binary | 0.98 | NA | [4] |
| litter_or_clutch_size_n | Clutch size | Continuous | 0.98 | NA | [3] |
| litters_or_clutches_per_y | Number of litters per year | Continuous | 0.85 | NA | [3] |
| log_area | Size of geographic distribution ($km^2$) | Continuous | 0.98 | $log_{10}$ | [1] |
| log_base_cort | Mean baseline corticosterone | Continuous | 0.26 | $log_{10}$ | [5] |
| log_birth_or_hatching_weight_g | Weight at hatching | Continuous | 0.44 | $log_{10}$ | [3] |
| log_ed_fair | Evolutionary distinctiveness (fair splits) | Continuous | 1 | $log_{10}$ | [6] |
| log_elevation_upper | Upper bounds of elevation across range | Continuous | 0.61 | $log_{10}$ | [4] |
| log_emass | Egg mass | Continuous | 0.93 | $log_{10}$ | [3] |

| | | | | | |
|---|---|---|---|---|---|
| log_femalemat | Days for female to reach sexual maturity | Continuous | 0.81 | $\log_{10}$ | [3] |
| log_fledging_age_d | Days until fledging | Continuous | 0.75 | $\log_{10}$ | [3] |
| log_fmass | Mass at fledging | Continuous | 0.23 | $\log_{10}$ | [3] |
| log_longevity | Maximum longevity | Continuous | 0.87 | $\log_{10}$ | [3] |
| log_malemat | Days for male to reach sexual maturity | Continuous | 0.75 | $\log_{10}$ | [3] |
| log_mass | Body mass of adults | Continuous | 1 | $\log_{10}$ | [3] |
| log_stress_cort | Mean stress-induced corticosterone | Continuous | 0.21 | $\log_{10}$ | [5] |
| marine_system | Does species inhabit marine ecosystem | Binary | 0.98 | NA | [4] |
| mig_disp | Size of non-breeding distribution relative to breeding distribution | Continuous | 0.98 | NA | [1] |
| mig_fullmigrant | Migratory strategy: full migrant | Binary | 0.98 | NA | [1] |
| mig_partialmigrant | Migratory strategy: partial migrant | Binary | 0.98 | NA | [1] |
| mig_resident | Migratory strategy: resident | Binary | 0.98 | NA | [1] |
| passerine | Is species in the Passeriformes | Binary | 0.98 | NA | [2,3] |
| pop_decreasing | IUCN population trend: decreasing | Binary | 0.98 | NA | [4] |
| pop_increasing | IUCN population trend: increasing | Binary | 0.98 | NA | [4] |
| pop_stable | IUCN population trend: stable | Binary | 0.98 | NA | [4] |
| ptaxa1 | First phylogenetic factor for competence | Binary | 1 | NA | [7] |
| s_mdist | Distance (km) between breeding and non-breeding distributions | Continuous | 0.98 | sqrt | [1] |
| swos | Web of Science citations | Continuous | 1 | sqrt | *rwos* |
| winter_lat | Latitude of the non-breeding distribution centroid | Continuous | 0.98 | NA | [1] |

*References*


1. BirdLife International, NatureServe. 2014 Bird species distribution maps of the world. *Cambridge, UK: BirdLife International*

2. Wilman H, Belmaker J, Simpson J, de la Rosa C, Rivadeneira MM, Jetz W. 2014 EltonTraits 1.0: Species-level foraging attributes of the world's birds and mammals. *Ecology* **95**, 2027–2027. (doi:10.1890/13-1917.1)

3. Myhrvold NP, Baldridge E, Chan B, Sivam D, Freeman DL, Ernest SKM. 2015 An amniote life-history database to perform comparative analyses with birds, mammals, and reptiles. *Ecology* **96**, 3109–3109. (doi:10.1890/15-0846R.1)

4. Chamberlain S. 2016 *rredlist:"IUCN" Red List Client. R package version 0.1. 0.*

5. Vitousek MN *et al.* 2018 HormoneBase, a population-level database of steroid hormone levels across vertebrates. *Scientific data* **5**, 180097.

6. Kembel SW, Cowan PD, Helmus MR, Cornwell WK, Morlon H, Ackerly DD, Blomberg SP, Webb CO. 2010 Picante: R tools for integrating phylogenies and ecology. *Bioinformatics* **26**, 1463–1464.

7. Washburne AD, Silverman JD, Morton JT, Becker DJ, Crowley D, Mukherjee S, David LA, Plowright RK. 2019 Phylofactorization: a graph partitioning algorithm to identify phylogenetic scales of ecological data. *Ecological Monographs* **0**, e01353. (doi:10.1002/ecm.1353)

### D. Phylogenetic meta-analysis

Table S2. Comparison of meta-analysis models for associations between space, year, and *Bb*sl prevalence of wild bird–derived tick larvae (*n*=777 after removing missing values). Models are ranked by ΔAICc with the number of coefficients (*k*), Akaike weights ($w_i$), and pseudo-$R^2$.

| **Fixed effects** | ***k*** | **ΔAICc** | $w_i$ | $R^2$ |
|---|---|---|---|---|
| ~ 1 | 1 | 0.00 | 0.48 | 0 |
| ~ year | 2 | 1.85 | 0.19 | 0 |
| ~ \|latitude\| | 2 | 1.96 | 0.18 | 0 |
| ~ year + \|latitude\|[1] | 3 | 3.82 | 0.07 | 0 |
| ~ UN georegion | 4 | 4.85 | 0.04 | 0 |
| ~ year + UN georegion[2] | 5 | 6.84 | 0.02 | 0 |
| ~ \|latitude\| + UN georegion | 5 | 6.86 | 0.02 | 0 |
| ~ year + UN georegion + \|latitude\|[3] | 6 | 8.87 | 0.01 | 0 |
| ~ \|latitude\| * UN georegion | 7 | 10.90 | <0.01 | 0 |
| ~ year + \|latitude\| * UN georegion | 8 | 12.92 | <0.01 | 0 |

[1]year * |latitude| was excluded owing to failed convergence
[2]year * UN georegion was excluded owing to failed convergence
[3]year * UN georegion + |latitude| was excluded owing to failed convergence

Table S3. Comparison of meta-analysis models for associations between seasons, space, and *Bb*sl prevalence of wild bird–derived tick larvae (*n*=604 after removing missing values). Models are ranked by ΔAICc with the number of coefficients (*k*), Akaike weights ($w_i$), and pseudo-$R^2$.

| **Fixed effects** | ***k*** | **ΔAICc** | $w_i$ | $R^2$ |
|---|---|---|---|---|
| ~ summer | 2 | 0.00 | 0.44 | 0.01 |
| ~ summer + \|latitude\| | 3 | 1.89 | 0.17 | 0 |
| ~ 1 | 1 | 3.43 | 0.08 | 0 |
| ~ summer * \|latitude\| | 4 | 3.8 | 0.07 | 0 |
| ~ winter | 2 | 4.72 | 0.04 | 0 |
| ~ fall | 2 | 5.04 | 0.04 | 0 |
| ~ spring | 2 | 5.45 | 0.03 | 0 |
| ~ summer + UN georegion | 5 | 5.63 | 0.03 | 0 |
| ~ winter * \|latitude\| | 4 | 6.27 | 0.02 | 0 |
| ~ spring * \|latitude\| | 4 | 6.64 | 0.02 | 0 |
| ~ winter + \|latitude\| | 3 | 6.76 | 0.02 | 0 |
| ~ fall + \|latitude\| | 3 | 7.07 | 0.01 | 0 |
| ~ summer * UN georegion | 6 | 7.43 | 0.01 | 0 |
| ~ spring + \|latitude\| | 3 | 7.48 | 0.01 | 0 |
| ~ spring * UN georegion | 6 | 8.68 | 0.01 | 0.02 |
| ~ fall * \|latitude\| | 4 | 8.75 | 0.01 | 0 |
| ~ winter + UN georegion | 5 | 10.18 | 0 | 0 |
| ~ fall + UN georegion | 5 | 10.67 | 0 | 0 |
| ~ spring + UN georegion | 5 | 11.17 | 0 | 0 |
| ~ winter * UN georegion | 6 | 11.57 | 0 | 0 |
| ~ fall * UN georegion | 6 | 12.08 | 0 | 0 |

### E. Phylogenetic factorization

Figure S3. Phylogenetic patterns in avian *Bb*sl competence (left) and study effort (Web of Science citation counts; right). Clades with significantly different competence or study effort from the paraphyletic remainder using phylogenetic factorization are highlighted.

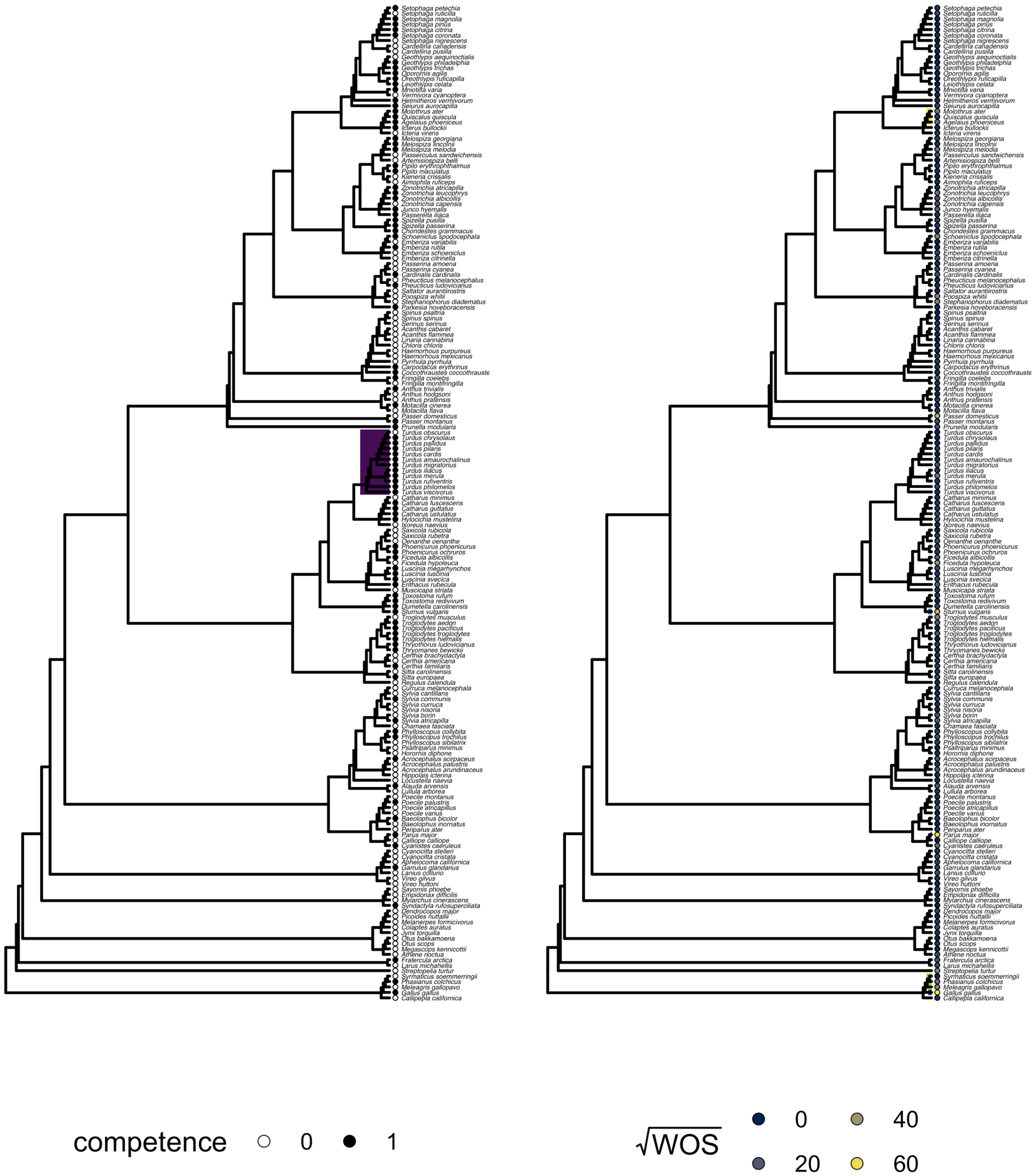

### F. Boosted regression trees

Table S4. Feature relative importance (mean and SE) of competence BRTs.

| Feature | Mean | SE |
|---|---|---|
| log_base_cort | 0.08 | 0.01 |
| log_fledging_age_d | 0.07 | 0.01 |
| log_longevity | 0.06 | 0 |
| mig_disp | 0.06 | 0 |
| breed_lat | 0.06 | 0 |
| log_elevation_upper | 0.06 | 0 |
| swos | 0.05 | 0 |
| log_area | 0.05 | 0 |
| log_birth_or_hatching_weight_g | 0.05 | 0 |
| log_fmass | 0.04 | 0 |
| log_emass | 0.04 | 0 |
| incubation_d | 0.04 | 0 |
| winter_lat | 0.03 | 0 |
| log_ed_fair | 0.03 | 0 |
| s_mdist | 0.03 | 0 |
| pop_decreasing | 0.02 | 0 |
| litter_or_clutch_size_n | 0.02 | 0 |
| log_femalemat | 0.02 | 0 |
| ptaxa1 | 0.02 | 0 |
| log_mass | 0.02 | 0 |
| litters_or_clutches_per_y | 0.02 | 0 |
| Diet.Seed | 0.01 | 0 |
| ForStrat.ground | 0.01 | 0 |
| passerine | 0.01 | 0 |
| ForStrat.midhigh | 0.01 | 0 |
| family_Turdidae | 0.01 | 0 |
| Diet.Inv | 0.01 | 0 |
| ForStrat.understory | 0.01 | 0 |
| Diet.Nect | 0.01 | 0 |
| diet_diversity | 0.01 | 0 |
| family_Fringillidae | 0.01 | 0 |
| family_Parulidae | 0.01 | 0 |
| ForStrat.canopy | 0.01 | 0 |
| Diet.PlantO | 0 | 0 |
| log_stress_cort | 0 | 0 |
| Diet.Fruit | 0 | 0 |
| pop_increasing | 0 | 0 |
| log_malemat | 0 | 0 |
| marine_system | 0 | 0 |
| pop_stable | 0 | 0 |

| | | |
|---|---|---|
| freshwater_system | 0 | 0 |
| family_Emberizidae | 0 | 0 |
| foraging_diversity | 0 | 0 |
| mig_fullmigrant | 0 | 0 |
| mig_partialmigrant | 0 | 0 |
| Diet.Vect | 0 | 0 |
| family_Sylviidae | 0 | 0 |
| Diet.Vend | 0 | 0 |
| family_Paridae | 0 | 0 |
| mig_resident | 0 | 0 |
| family_Muscicapidae | 0 | 0 |
| IUCN_LC | 0 | 0 |
| Diet.Scav | 0 | 0 |

Table S5. Phylogenetic factorization of (logit-transformed) mean predicted probabilities for *Bb*sl competence across the 39 avian families. Shown are the number of retained clades after applying Holm's sequentially rejective 5% cutoff for the family-wise error rate, taxa corresponding to those clades, number of species per clade, and mean probabilities for the clade compared to the paraphyletic remainder. Clades with higher predicted probabilities are displayed in grey.

| **Clade** | **Taxa** | **Species** | **Clade** | **Other** |
|---|---|---|---|---|
| 1 | *Turdus, Nesocichla, Psophocichla* | 79 | 0.49 | 0.33 |
| 2 | *Chlorospingus, Rhynchospiza, Aimophila, Arremonops, Ammodramus, Peucaea, Melozone, Kieneria, Torreornis, Pezopetes, Pipilo, Pselliophorus, Atlapetes, Melospiza, Xenospiza, Passerculus, Pooecetes, Oriturus, Artemisiospiza, Zonotrichia, Junco, Spizella, Passerella, Arremon, Amphispiza, Chondestes, Calamospiza, Emberiza, Schoeniclus, Latoucheornis, Melophus, Myiothlypis, Myioborus, Cardellina, Basileuterus, Dendroica, Setophaga, Parula, Catharopeza, Geothlypis, Oporornis, Leiothlypis, Vermivora, Oreothlypis, Protonotaria, Limnothlypis, Mniotilta, Helmitheros, Seiurus,* Icteridae*, Teretistris, Zeledonia, Icteria, Spindalis, Nesospingus, Microligea, Phaenicophilus, Calyptophilus* | 373 | 0.38 | 0.33 |
| 3 | *Zonotrichia leucophrys, Zonotrichia atricapilla, Zonotrichia albicollis, Zonotrichia querula* | 4 | 0.78 | 0.33 |
| 4 | Strigidae, Picidae | 428 | 0.29 | 0.34 |
| 5 | Alcidae | 25 | 0.47 | 0.33 |
| 6 | Mimidae | 34 | 0.42 | 0.33 |
| 7 | Parulidae | 2 | 0.72 | 0.33 |
| 8 | *Sylvia atricapilla* | 1 | 0.83 | 0.33 |
| 9 | *Geokichla, Zoothera, Cochoa, Catharus, Entomodestes, Cichlopsis, Hylocichla, Ixoreus, Sialia, Myadestes townsendi, Myadestes obscurus, Myadestes unicolor, Myadestes occidentalis, Myadestes coloratus, Myadestes melanops, Myadestes ralloides, Myadestes genibarbis, Myadestes elisabeth, Myadestes lanaiensis, Myadestes myadestinus, Neocossyphus, Stizorhina, Grandala* | 76 | 0.38 | 0.33 |

Table S6. Mean predicted probabilities of avian competence for *Bb*sl from the BRTs. Species with propensity scores above 50% are shown, with those above 60% shaded in grey.

| Latin binomial | Common name | *P* | Competence |
|---|---|---|---|
| *Sylvia atricapilla* | Eurasian Blackcap | 0.827 | positive |
| *Zonotrichia albicollis* | White-throated Sparrow | 0.816 | positive |
| *Zonotrichia leucophrys* | White-crowned Sparrow | 0.814 | positive |
| *Pipilo erythrophthalmus* | Eastern Towhee | 0.808 | positive |
| *Turdus merula* | Common Blackbird | 0.804 | positive |
| *Leiothlypis celata* | Orange-crowned Warbler | 0.802 | positive |
| *Molothrus ater* | Brown-headed Cowbird | 0.8 | positive |
| *Turdus migratorius* | American Robin | 0.784 | positive |
| *Luscinia svecica* | Bluethroat | 0.784 | positive |
| *Melospiza melodia* | Song Sparrow | 0.784 | positive |
| *Parkesia noveboracensis* | Northern Waterthrush | 0.784 | positive |
| *Zonotrichia atricapilla* | Golden-crowned Sparrow | 0.775 | positive |
| *Quiscalus quiscula* | Common Grackle | 0.77 | positive |
| *Cyanistes caeruleus* | Eurasian Blue Tit | 0.76 | positive |
| *Catharus guttatus* | Hermit Thrush | 0.757 | positive |
| *Spinus tristis* | American Goldfinch | 0.752 | unsampled |
| *Catharus ustulatus* | Swainson's Thrush | 0.749 | positive |
| *Agelaius phoeniceus* | Red-winged Blackbird | 0.747 | positive |
| *Fringilla coelebs* | Common Chaffinch | 0.746 | positive |
| *Spizella passerina* | Chipping Sparrow | 0.736 | positive |
| *Hylocichla mustelina* | Wood Thrush | 0.731 | positive |
| *Erithacus rubecula* | European Robin | 0.726 | positive |
| *Sturnus vulgaris* | Common Starling | 0.724 | positive |
| *Turdus philomelos* | Song Thrush | 0.723 | positive |
| *Phoenicurus phoenicurus* | Common Redstart | 0.722 | positive |
| *Melospiza georgiana* | Swamp Sparrow | 0.721 | positive |
| *Junco hyemalis* | Dark-eyed Junco | 0.719 | positive |
| *Turdus pilaris* | Fieldfare | 0.718 | positive |
| *Dumetella carolinensis* | Grey Catbird | 0.717 | positive |
| *Zonotrichia querula* | Harris's Sparrow | 0.716 | unsampled |
| *Seiurus aurocapilla* | Ovenbird | 0.711 | positive |
| *Parus major* | Great Tit | 0.709 | positive |
| *Uria lomvia* | Thick-billed Murre | 0.709 | unsampled |
| *Geothlypis trichas* | Common Yellowthroat | 0.709 | positive |
| *Eremophila alpestris* | Horned Lark | 0.707 | unsampled |
| *Melozone aberti* | Abert's Towhee | 0.707 | unsampled |
| *Turdus viscivorus* | Mistle Thrush | 0.701 | positive |
| *Xanthocephalus xanthocephalus* | Yellow-headed Blackbird | 0.698 | unsampled |
| *Setophaga coronata* | Myrtle Warbler | 0.697 | positive |
| *Toxostoma rufum* | Brown Thrasher | 0.696 | positive |
| *Passerella iliaca* | Fox Sparrow | 0.694 | positive |
| *Setophaga townsendi* | Townsend's Warbler | 0.69 | unsampled |
| *Piranga olivacea* | Scarlet Tanager | 0.689 | unsampled |
| *Phoenicurus ochruros* | Black Redstart | 0.689 | positive |
| *Troglodytes aedon* | House Wren | 0.683 | positive |

| | | | |
|---|---|---|---|
| *Pheucticus ludovicianus* | Rose-breasted Grosbeak | 0.681 | positive |
| *Melospiza lincolnii* | Lincoln's Sparrow | 0.679 | positive |
| *Prunella modularis* | Dunnock | 0.677 | positive |
| *Sialia sialis* | Eastern Bluebird | 0.67 | unsampled |
| *Setophaga petechia* | Mangrove Warbler | 0.67 | positive |
| *Spizella pusilla* | Field Sparrow | 0.669 | positive |
| *Troglodytes troglodytes* | Eurasian Wren | 0.668 | positive |
| *Sturnella neglecta* | Western Meadowlark | 0.663 | unsampled |
| *Sitta europaea* | Eurasian Nuthatch | 0.662 | positive |
| *Mimus polyglottos* | Northern Mockingbird | 0.661 | unsampled |
| *Parkesia motacilla* | Louisiana Waterthrush | 0.651 | unsampled |
| *Euphagus cyanocephalus* | Brewer's Blackbird | 0.648 | unsampled |
| *Cardinalis cardinalis* | Northern Cardinal | 0.648 | positive |
| *Pooecetes gramineus* | Vesper Sparrow | 0.647 | unsampled |
| *Turdus iliacus* | Redwing | 0.646 | positive |
| *Luscinia megarhynchos* | Common Nightingale | 0.639 | positive |
| *Pipilo maculatus* | Spotted Towhee | 0.638 | positive |
| *Oreothlypis ruficapilla* | Nashville Warbler | 0.637 | positive |
| *Emberiza calandra* | Corn Bunting | 0.628 | unsampled |
| *Cerorhinca monocerata* | Rhinoceros Auklet | 0.627 | unsampled |
| *Vireo olivaceus* | Red-eyed Vireo | 0.626 | unsampled |
| *Setophaga magnolia* | Magnolia Warbler | 0.623 | positive |
| *Ammodramus savannarum* | Grasshopper Sparrow | 0.621 | unsampled |
| *Phylloscopus collybita* | Common Chiffchaff | 0.62 | positive |
| *Helmitheros vermivorum* | Worm-eating Warbler | 0.615 | positive |
| *Empidonax virescens* | Acadian Flycatcher | 0.614 | unsampled |
| *Catharus fuscescens* | Veery | 0.613 | positive |
| *Turdus grayi* | Clay-colored Thrush | 0.611 | unsampled |
| *Passerina cyanea* | Indigo Bunting | 0.609 | negative |
| *Troglodytes pacificus* | Pacific Wren | 0.606 | positive |
| *Turdus amaurochalinus* | Creamy-bellied Thrush | 0.606 | positive |
| *Troglodytes hiemalis* | Winter Wren | 0.603 | positive |
| *Chondestes grammacus* | Lark Sparrow | 0.601 | positive |
| *Sylvia communis* | Common Whitethroat | 0.597 | positive |
| *Sterna hirundo* | Common Tern | 0.595 | unsampled |
| *Uria aalge* | Common Murre | 0.593 | unsampled |
| *Icterus galbula* | Baltimore Oriole | 0.591 | unsampled |
| *Spinus pinus* | Pine Siskin | 0.583 | unsampled |
| *Icterus bullockii* | Bullock's Oriole | 0.582 | positive |
| *Oreoscoptes montanus* | Sage Thrasher | 0.581 | unsampled |
| *Sialia currucoides* | Mountain Bluebird | 0.581 | unsampled |
| *Setophaga citrina* | Hooded Warbler | 0.581 | positive |
| *Turdus rufiventris* | Rufous-bellied Thrush | 0.581 | positive |
| *Turdus cardis* | Japanese Thrush | 0.581 | positive |
| *Emberiza pallasi* | Pallas's Reed Bunting | 0.58 | unsampled |
| *Acrocephalus scirpaceus* | Eurasian Reed Warbler | 0.579 | positive |
| *Turdus subalaris* | Eastern Slaty Thrush | 0.578 | unsampled |
| *Alauda arvensis* | Eurasian Skylark | 0.577 | positive |
| *Toxostoma bendirei* | Bendire's Thrasher | 0.577 | unsampled |

| | | | |
|---|---|---|---|
| *Corvus brachyrhynchos* | American Crow | 0.575 | unsampled |
| *Turdus hortulorum* | Grey-backed Thrush | 0.574 | unsampled |
| *Coccothraustes coccothraustes* | Hawfinch | 0.574 | positive |
| *Ficedula albicollis* | Collared Flycatcher | 0.573 | positive |
| *Turdus torquatus* | Ring Ouzel | 0.572 | unsampled |
| *Catharus bicknelli* | Bicknell's Thrush | 0.572 | unsampled |
| *Icterus spurius* | Orchard Oriole | 0.572 | unsampled |
| *Turdus obscurus* | Eyebrowed Thrush | 0.572 | negative |
| *Geothlypis philadelphia* | Mourning Warbler | 0.568 | positive |
| *Empidonax alnorum* | Alder Flycatcher | 0.567 | unsampled |
| *Phasianus colchicus* | Common Pheasant | 0.567 | positive |
| *Pica pica* | Eurasian Magpie | 0.567 | unsampled |
| *Geothlypis tolmiei* | MacGillivray's Warbler | 0.566 | unsampled |
| *Fratercula arctica* | Atlantic Puffin | 0.565 | positive |
| *Lanius ludovicianus* | Loggerhead Shrike | 0.564 | unsampled |
| *Vermivora chrysoptera* | Golden-winged Warbler | 0.564 | unsampled |
| *Luscinia luscinia* | Thrush Nightingale | 0.564 | positive |
| *Passer montanus* | Eurasian Tree Sparrow | 0.562 | positive |
| *Turdus pallidus* | Pale Thrush | 0.562 | positive |
| *Thryothorus ludovicianus* | Carolina Wren | 0.562 | positive |
| *Mniotilta varia* | Black-and-white Warbler | 0.555 | positive |
| *Myiarchus crinitus* | Great Crested Flycatcher | 0.555 | unsampled |
| *Turdus nigriceps* | Andean Slaty Thrush | 0.554 | unsampled |
| *Setophaga caerulescens* | Black-throated Blue Warbler | 0.553 | unsampled |
| *Serinus canaria* | Atlantic Canary | 0.553 | unsampled |
| *Setophaga fusca* | Blackburnian Warbler | 0.552 | unsampled |
| *Rissa tridactyla* | Black-legged Kittiwake | 0.552 | unsampled |
| *Setophaga pinus* | Pine Warbler | 0.551 | positive |
| *Passerina ciris* | Painted Bunting | 0.55 | unsampled |
| *Euphagus carolinus* | Rusty Blackbird | 0.55 | unsampled |
| *Setophaga kirtlandii* | Kirtland's Warbler | 0.55 | unsampled |
| *Sturnella magna* | Eastern Meadowlark | 0.547 | unsampled |
| *Alle alle* | Little Auk | 0.547 | unsampled |
| *Dolichonyx oryzivorus* | Bobolink | 0.546 | unsampled |
| *Spizella arborea* | American Tree Sparrow | 0.543 | unsampled |
| *Setophaga auduboni* | Audubon's Warbler | 0.541 | unsampled |
| *Turdus lherminieri* | Forest Thrush | 0.538 | unsampled |
| *Turdus leucomelas* | Pale-breasted Thrush | 0.536 | unsampled |
| *Emberiza schoeniclus* | Common Reed Bunting | 0.535 | negative |
| *Turdus simillimus* | Indian Blackbird | 0.534 | unsampled |
| *Spiza americana* | Dickcissel | 0.534 | unsampled |
| *Toxostoma longirostre* | Long-billed Thrasher | 0.534 | unsampled |
| *Geospiza fuliginosa* | Small Ground Finch | 0.533 | unsampled |
| *Lagopus lagopus* | Willow Ptarmigan | 0.531 | unsampled |
| *Geothlypis formosa* | Kentucky Warbler | 0.529 | unsampled |
| *Larus delawarensis* | Ring-billed Gull | 0.528 | unsampled |
| *Icteria virens* | Yellow-breasted Chat | 0.528 | negative |
| *Aphelocoma coerulescens* | Florida Scrub Jay | 0.527 | unsampled |
| *Turdus aurantius* | White-chinned Thrush | 0.526 | unsampled |

| | | | |
|---|---|---|---|
| *Alca torda* | Razorbill | 0.526 | unsampled |
| *Agelaius tricolor* | Tricolored Blackbird | 0.526 | unsampled |
| *Phylloscopus trochilus* | Willow Warbler | 0.526 | positive |
| *Gallus gallus* | Red Junglefowl | 0.525 | positive |
| *Acrocephalus dumetorum* | Blyth's Reed Warbler | 0.525 | unsampled |
| *Leucosticte tephrocotis* | Grey-crowned Rosy Finch | 0.525 | unsampled |
| *Aethia cristatella* | Crested Auklet | 0.525 | unsampled |
| *Turdus ruficollis* | Red-throated Thrush | 0.523 | unsampled |
| *Locustella luscinioides* | Savi's Warbler | 0.52 | unsampled |
| *Turdus lawrencii* | Lawrence's Thrush | 0.52 | unsampled |
| *Vireo solitarius* | Blue-headed Vireo | 0.519 | unsampled |
| *Protonotaria citrea* | Prothonotary Warbler | 0.519 | unsampled |
| *Turdus falcklandii* | Austral Thrush | 0.518 | unsampled |
| *Thryomanes bewickii* | Bewick's Wren | 0.518 | positive |
| *Larus glaucescens* | Glaucous-winged Gull | 0.518 | unsampled |
| *Pipilo chlorurus* | Green-tailed Towhee | 0.518 | unsampled |
| *Garrulus glandarius* | Eurasian Jay | 0.518 | positive |
| *Setophaga pensylvanica* | Chestnut-sided Warbler | 0.517 | unsampled |
| *Zenaida macroura* | Mourning Dove | 0.516 | unsampled |
| *Coturnix japonica* | Japanese Quail | 0.516 | unsampled |
| *Geospiza scandens* | Common Cactus Finch | 0.515 | unsampled |
| *Sialia mexicana* | Western Bluebird | 0.514 | unsampled |
| *Turdus jamaicensis* | White-eyed Thrush | 0.514 | unsampled |
| *Turdus eunomus* | Dusky Thrush | 0.513 | unsampled |
| *Turdus naumanni* | Naumann's Thrush | 0.512 | unsampled |
| *Turdus chrysolaus* | Brown-headed Thrush | 0.511 | positive |
| *Cepphus columba* | Pigeon Guillemot | 0.51 | unsampled |
| *Turdus nudigenis* | Spectacled Thrush | 0.51 | unsampled |
| *Turdus swalesi* | La Selle Thrush | 0.509 | unsampled |
| *Coturnix coturnix* | Common Quail | 0.505 | unsampled |
| *Motacilla alba* | White Wagtail | 0.503 | unsampled |
| *Piranga rubra* | Summer Tanager | 0.503 | unsampled |
| *Vireo griseus* | White-eyed Vireo | 0.501 | unsampled |
| *Larus argentatus* | European Herring Gull | 0.501 | unsampled |
| *Aegolius funereus* | Boreal Owl | 0.501 | unsampled |

Figure S4. Distribution of the mean predicted probabilities of *Bb*sl competence across the 39 sampled avian families. (A) Density plots show predictions for currently negative, positive, and unsampled species, and these propensity scores are also shown across the avian phylogeny (B). Distributions of species within a mean probability of over 50% are shown by their breeding (C), non-breeding (D), and resident ranges (E).

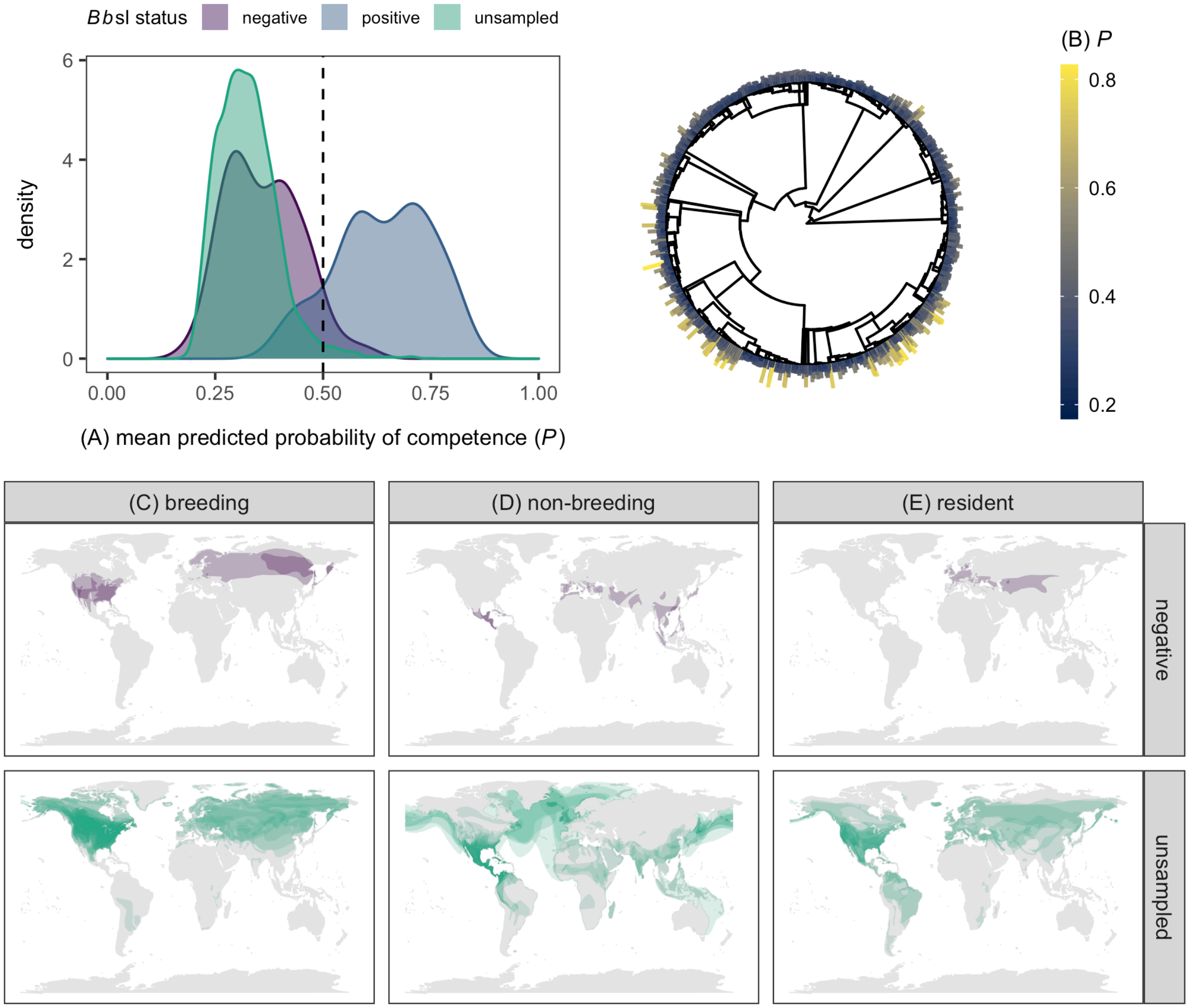
